# Optic cup folding is driven by the geometry and tension of the Retinal Pigmented Epithelium (RPE) cells

**DOI:** 10.64898/2026.07.03.736268

**Authors:** Juan Aperador-Redondo, Estefanía Sanabria-Reinoso, Javier Macho-Rendón, Rocío Polvillo, JR Martínez Morales, María Almuedo-Castillo

## Abstract

While optic cup folding is known to involve specific geometrical changes of RPE cells, the precise gene regulatory mechanisms orchestrating the adoption of their highly rigid geometry, and how these contribute to successful folding, remain poorly understood. To address this gap, we investigated how the increase in mechanical tension and maintenance of an elongated geometry depend on the activation of the Wnt/β-catenin and YAP pathways in RPE cells. We demonstrated that interference with these pathways causes folding failure due to a reduction in RPE cellular tension. We also identified transcriptional programs controlled by these pathways that regulate the mechanical properties of the actin cytoskeleton, cell-to-cell and cell-to-ECM adhesions, and endocytosis. Finally, we hypothesized that the LINC complex, which transmits tension between the cell and nuclear membranes, is responsible for the nuclear entry of β-catenin and YAP in a cellular geometry-dependent manner.

We combined quantitative imaging, functional analysis, mechanical perturbation assays, and transcriptomic analysis to generate a comprehensive view of how the coordination of mechanosensitive gene expression and changes in cellular geometries drive eye formation.

## Introduction

How an individual epithelial sheet dynamically changes its mechanical properties and cellular geometries while buffering tissue-level forces to become a complex organ remains a fundamental question in developmental mechanobiology. During vertebrate eye formation, this question is nicely illustrated by the Retinal Pigmented Epithelium (RPE), the outer layer to the Neural retina (NR), which must rapidly transition from a rounded shape to a highly flattened and stretched geometry, a morphogenetic program required for optic cup folding (Moreno-Mármol *et al*, 2021). This complex folding process is driven by collective contributions from distinct eye subpopulations. While the NR undergoes basal constriction to generate inward bending forces (Nicolás-Pérez *et al*, 2016; Sidhaye & Norden, 2017), RPE cells experience strong lateral strain as they acquire their characteristic stretched and rigid epithelial configuration (Moreno-Mármol *et al*, 2021). Despite this profound cellular transformation, the precise mechanotransduction pathways that dictate this particular geometry and maintains tissue-level tension remains poorly understood.

To address these unanswered questions of tissue morphogenesis, we investigated how developing RPE cells actively sense, execute and maintain their unique mechanical properties. Previous studies have shown that the activation of Yes-associated protein (YAP), a primary mechanostransduction effector, is highly regionalized in the zebrafish eye and restricted to the RPE cells (Miesfeld *et al*, 2015). While homozygous *yap1* mutants display relatively milder ocular phenotypes, double mutants of *yap1* and its paralog *wwrt1* (*taz*) results in the arrestment of global development hours before the folding of the optic cup takes place (Kimelman *et al*, 2017; Miesfeld *et al*, 2015).

Here, we first observed that alongside YAP, the activation of Wnt/β-catenin pathway, involved in the specification of RPE cellular identity in the mouse optic cup (Westenskow *et al*, 2009), it is also restricted to the RPE cells in a sequential and transient manner. Moreover, our findings demonstrate that the coordinated activation of YAP and β-catenin signaling pathways is required to sustain the mechanical and geometrical signature of the RPE cells, and is consequently essential for the success of optic cup morphogenesis. Using a combination of *in vivo* quantitative imaging, primary zebrafish cell culture, laser-mediated mechanical ablations and transcriptomic analysis, we demonstrated that these two transcription factors are selectively activated in the RPE cells in response to high endogenous intracellular tension of the RPE tissue. Once active, they respond by maintaining this mechanical state, establishing a self-sustaining positive feedback loop to preserve and promote RPE cellular stretching. Pharmacological and genetic perturbations inhibiting YAP and β-catenin signaling revealed that delayed optic cup folding is directly explained by a strong reduction in actomyosin-mediated tissue tension at the RPE-NR interface and of the RPE cells, causing them to remain in an incomplete and rounded geometrical state.

Through comparative transcriptomic profiling, we identified a convergent gene regulatory program driven downstream of YAP, β-catenin, and the induction of actomyosin intracellular contractility. This shared transcriptional program co-regulates essential mechanical components of the cell, such as the actomyosin cytoskeleton, cell-to-cell and cell-to-ECM adhesions, and probably the force-sensing ion channel Piezo1. Furthermore, our data uncovers the possible involvement of the endocytic machinery to buffer membrane tension to accommodate the rapid membrane expansion experienced by stretching RPE cells.

Finally, using mosaic transplantation assays, we identify the Linker of Nucleoskeleton and Cytoskeleton complex (LINC) as the possible physical transducer linking the actomyosin tension to the nuclear translocation of YAP and β-catenin transcription factors to activate the mechanical gene expression program.

Altogether, our data integrates tension transducers (LINC complex), cytoskeletal anchors (actomyosin complex and cellular adhesions) and nuclear effectors (YAP and β-catenin) to provide a comprehensive, self-sustaining mechanochemical model required for the correct formation of the vertebrate eye.

## Results

### RPE-specific activity of YAP and β-catenin is essential for cell elongation and optic cup folding

The activation of YAP specifically at the RPE of developing zebrafish eye cups has been previously reported using a transgenic sensor for YAP/TEAD activation (*4XGTIIC:Kaede* or *TEAD:Kaede*) (Miesfeld & Link, 2014; Miesfeld *et al*, 2015; Voltes *et al*, 2019) (Figure 1B). Here, we found that β-catenin, another key mechanosensitive transcription factor (Cha *et al*, 2016; Ray *et al*, 2013), is also specifically activated in these RPE cells. Notably, the activation profile of β-catenin differed from that of YAP, as it occurred in a transient manner (Figure 1A and 1B; Supp video 1, 2, and 3). This dynamic nuclear translocation was probably resolvable due to the temporal sensitivity of the β-catenin reporter line (*7xTCFsiam:mCherry or TCF:mCherry*; (Moro *et al*, 2013), which allowed us to capture the transient nature of the signal across the RPE population (Supp videos 1, 2, and 3). The localized activation of the YAP and β-catenin pathways is further corroborated by single-cell expression data (Macho-Rendón *et al*, 2026). UMAP analysis revealed that downstream targets and effectors of both the Wnt (tcf12 and wnt2) and YAP (ctgfa and yap1) pathways were significantly enriched in clusters corresponding to RPE cells compared to the neural retina, supporting the tissue-specific roles of these transcription factors (Figure 1C).

**Figure 1.**
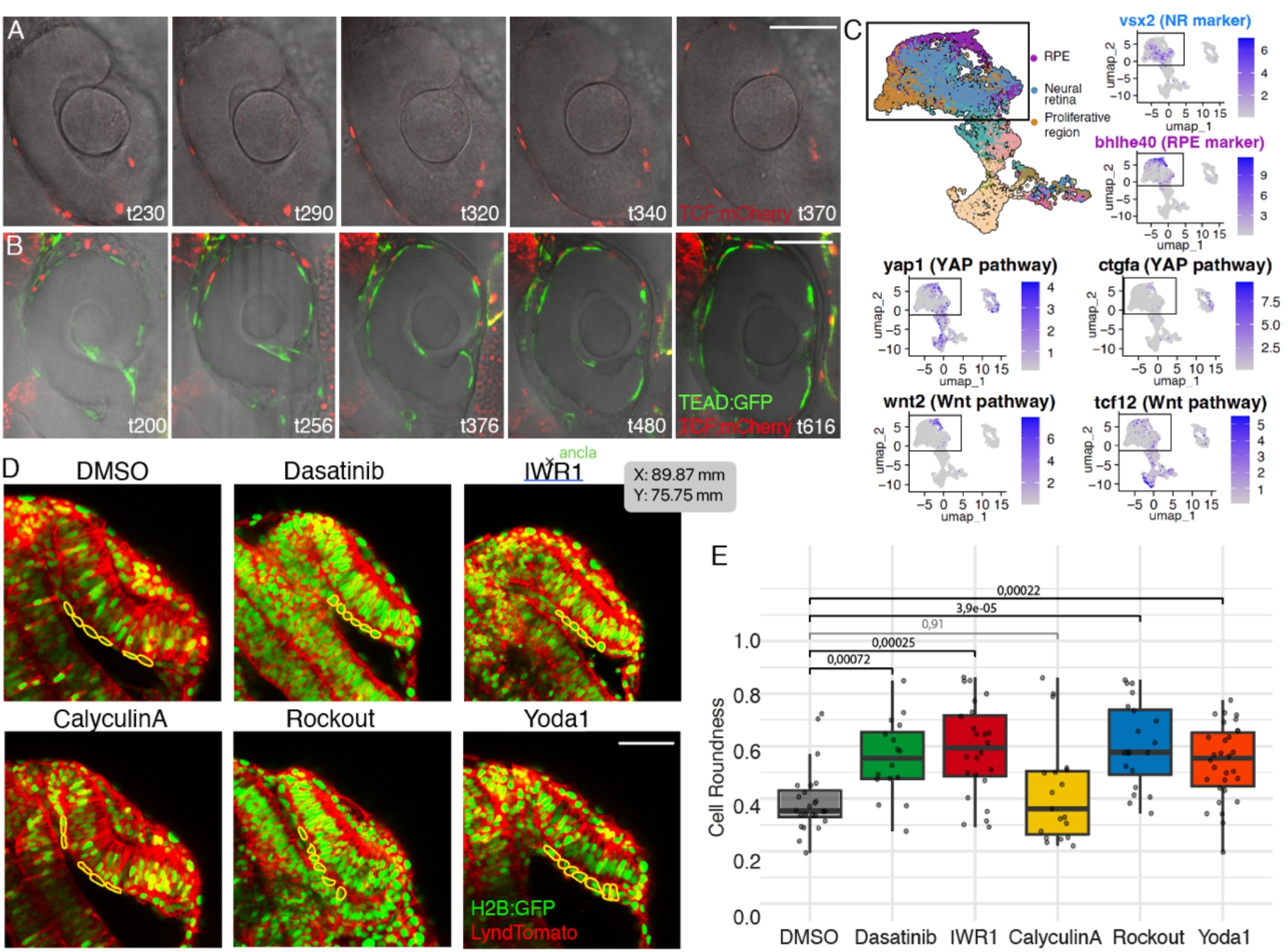
Specific activation of YAP and β-catenin signaling influences RPE cell geometry during optic cup folding. **(A)** Time-lapse snapshots of live imaging showing the spatiotemporal dynamics of Wnt/β-catenin activation in the RPE cells via the β-catenin reporter 7x*TCFsiam:*mCherry (*TCF:*mCherry) (red). Wnt/β-catenin is activated exclusively in the RPE cells in a sequential transient manner. These images correspond to Supplementary video 1. Time points (t) are indicated in minutes. **(B)** Time-lapse snapshots of live imaging showing the spatiotemporal dynamics of YAP and β-catenin activation in the RPE cells via the double reporter line for YAP 4X*GTIIC:*Kaede (*TEAD:*Kaede) (green) and β-catenin 7x*TCFsiam:*mCherry (*TCF:*mCherry) (red). YAP and β-catenin activity are co-localized and restricted to the flattening RPE domain. These images correspond to Supplementary video 3. Time points (t) are indicated in minutes. Note that t0 correspond to the initiation of time-lapse imaging for each respective movie and does not reflect an identical developmental stage between individual imaging sessions in (A) and (B). **(C)** UMAP clustering and scRNA-seq expression profiles of early zebrafish eye cells (Macho-Rendón *et al*, 2026). Clusters are segregated into RPE (purple, marked by *bhlhe40*), Neural Retina (blue, marked by *vsx2*), and proliferative regions (orange). Feature plots display the restricted and overlapping expression of YAP pathway effectors (*yap1*, *ctgfa*), and Wnt/β-catenin pathway components (*wnt2*, *tcf12*) within the RPE cluster. **(D)** Representative confocal images when the optic cup folds (22 hpf) in embryos expressing *Lyn-td*Tomato (membrane marker, red) and *h2B:*GFP (nuclear marker, green) following pharmacological treatments: Control with DMSO, Dasatinib (YAP inhibitor), IWR-1 (Wnt/β-catenin inhibitor), Calyculin A (increases myosin II phosphorylation and actomyosin contractility), Rockout (actomyosin contractility inhibitor) and Yoda1 (agonist for the mechanosensitive ion channel Piezo1). In yellow, representative RPE geometries are highlighted. **(E)** Quantification of RPE cell roundness across the different pharmacological treatments shown in (D). Box plots represent the distribution of cell roundness values; individual data points represent single cells. Data normality was assessed via the Shapiro-Wilk test. Because at least one group exhibited a non-normal distribution, statistical significance was determined using the non-parametric Wilcoxon test. n=24 DMSO cells from 3 embryos; n=16 Dasatinib-treated cells from 3 embryos; n=24 IWR1-treated cells from 3 embryos; n=19 CalyculinA-treated cells from 3 embryos; n=20 Rockout-treated cells from 3 embryos; n=30 Yoda1-treated cells from 3 embryos. Scale bars: 50 µm.

To explore the functional roles of YAP and β-catenin during RPE morphogenesis, we employed a pharmacological perturbation strategy. This approach allowed for precise dose-dependent and temporal control, which was essential for bypassing earlier developmental requirements for these pathways, such as gastrulation and posterior body elongation. By optimizing drug concentrations, we achieved robust inhibition of optic cup folding without compromising overall embryonic viability (Supp Figure 1). Among several candidates, we selected Dasatinib (a YAP inhibitor; (Sousa-Ortega *et al*, 2023) and IWR-1 (a Wnt/β-catenin inhibitor; (Chen *et al*, 2009) for their higher specificity in the eye tissue. Furthermore, to investigate the interplay between these transcription factors and mechanical forces, we modulated intracellular tension using Rockout (an inhibitor of actomyosin contractility;(Sidhaye & Norden, 2017), Calyculin A (which increases myosin II phosphorylation and actomyosin contractility; (Ishihara *et al*, 1989), and Yoda1 (an agonist for the mechanosensitive ion channel Piezo1; (Syeda *et al*, 2015). Piezo1 is the most known mechanosensitive ion channel that convert physical forces into chemical signals. Yoda1 is an small-molecule agonist that results in the hyperactivation of these mechanosensitive channels and leads to an aberrant channel gating that unable cells to accurately perceive environmental cues, rather than producing an overall increase in functional tissue tension (Botello-Smith *et al*, 2019). Interestingly, we observed a consistent and significant delay in optic cup folding of about 4-5 hours, following the pharmacological inhibition of YAP (Dasatinib), β-catenin (IWR-1), Rho-kinase-mediated tension (Rockout), and mechanosensation (Yoda1) (Figure 1D and Supp Figure 1). In contrast, promoting the opposite effect, which is to enhance intracellular tension via Calyculin A, did not alter the folding kinetics (Figure 1D and Supp Figure 1). Similarly, this delay was accompanied by a lack of stretching of the RPE cells when the ability to transduce tension was pharmacologically inhibited, but not when the intracellular tension was increased (Figure 1E). These results support the idea that RPE cells operate under a high basal level of intracellular tension (Moreno-Mármol et al., 2021), such that further increases do not alter cellular geometries or accelerate morphogenesis, whereas any reduction in this mechanical tension might significantly impair the folding process.

To validate the specificity of the YAP and β-catenin pharmacological inhibitors during eye morphogenesis, we employed alternative genetic strategies to perturb these signaling pathways. First, we injected mRNA encoding a dominant-negative form of YAP (*yap*DN) at a titrated concentration that delayed the kinetics of optic cup folding without compromising overall embryonic development. Second, we generated a CRISPR-KO line for *wnt2*, which was identified as the most RPE-specific Wnt ligand in a recent single-cell transcriptomic analysis of the developing zebrafish eye. These genetic approaches recapitulated the phenotypes observed with pharmacological inhibition, which is a consistent delay in optic cup folding accompanied by RPE cells that maintained rounded, rather than flattened, cellular geometries (Supp Figure 2). To assess the functional impact of these changes on tissue mechanics, we analyzed the degree of actin filament contractility over time by monitoring Utrophin accumulation in the RPE of *YapDN* and *wnt2* KO embryos. Utrophin specifically labels stable, tension-loaded F-actin, serving as a proxy for contractile force. In our *in vivo* imaging experiments, we observed a marked reduction in the Utrophin signal, suggesting lower overall contractility (Supp Videos 4-7). Consequently, RPE cells in these embryos exhibit incomplete flattening and significant optic cup folding delays (Supp Figure 2).

**Figure 2.**
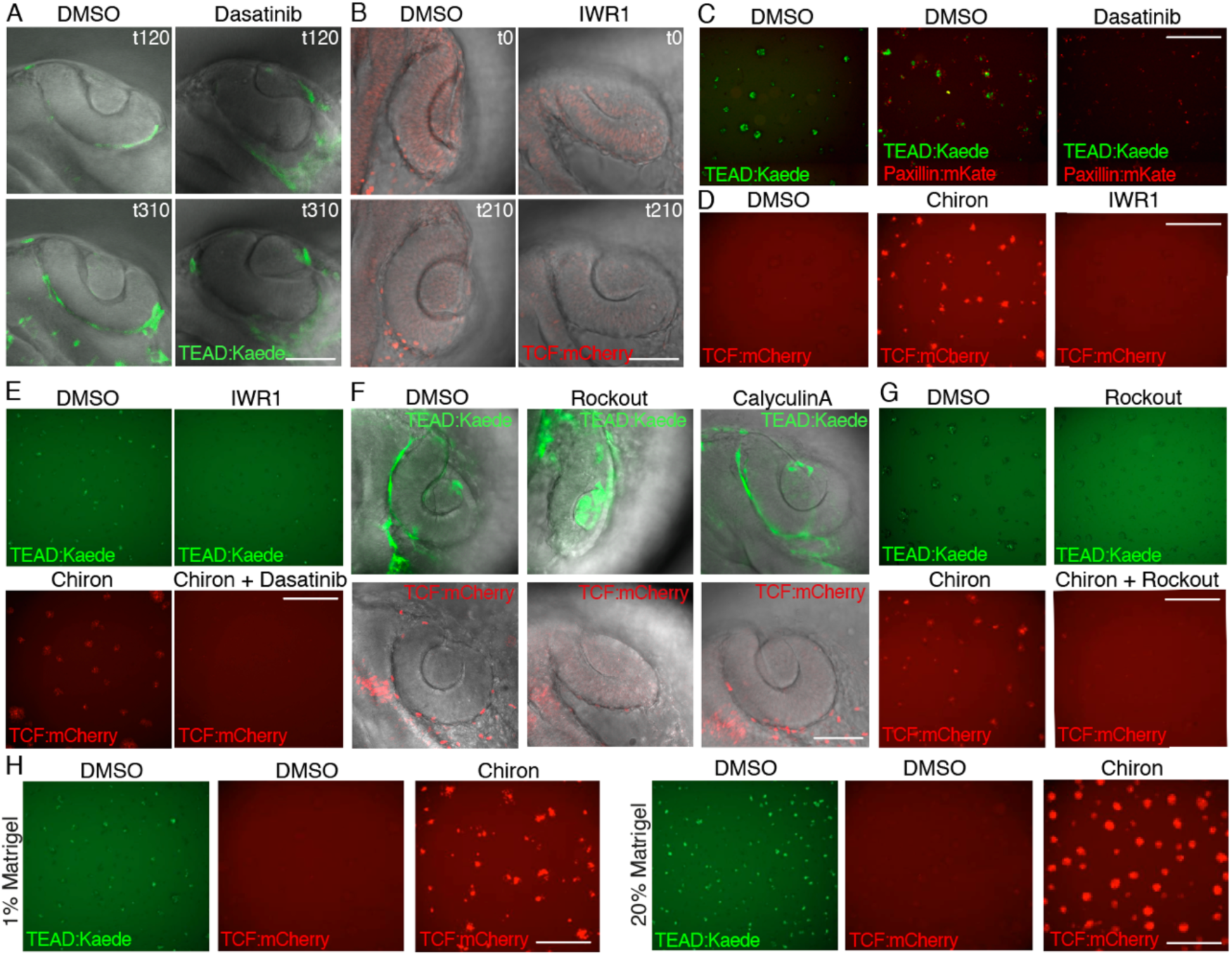
Mechanosensitive regulation of YAP and β-catenin signaling *in vivo* and *in vitro*. **(A)** Time-lapse live imaging showing the spatiotemporal dynamics of YAP activation in the RPE cells via the YAP reporter 4X*GTIIC:*Kaede (*TEAD:*Kaede) (green) in DMSO-treated (Control) and Dasatinib-treated (YAP inhibitor) embryos. Even when YAP signaling is just mildly inhibited using low dosis of Dasatininb, the YAP reporter *TEAD*:Kaede does not show activity in the RPE, validating the pharmacological treatment. These images correspond to Supplementary videos 8 and 9. Time points (t) are indicated in minutes. Scale bars: 50 µm. **(B)** Time-lapse live imaging showing the spatiotemporal dynamics of Wnt/β-catenin activation in the RPE cells via the β-catenin reporter 7x*TCFsiam:*mCherry (*TCF:*mCherry) (red) in DMSO-treated (Control) and IWR1-treated (β-catenin inhibitor) embryos. Even when β-catenin signaling is just mildly inhibited using low-dose of IWR1, the Wnt/β-catenin reporter *TCF:*mCherry does not show activity in the RPE, validating the pharmacological treatment. These images correspond to Supplementary videos 10 and 11. Time points (t) are indicated in minutes. Note that t0 correspond to the initiation of time-lapse imaging for each respective movie and does not reflect an identical developmental stage between individual imaging sessions in (A) and (B). Scale bars: 50 µm. **(C)** Still images from live imaging of dissociated zebrafish blastula cells from the reporter *TEAD:*Kaede (green) cultured in DMSO (Control) and Dasatinib. The focal adhesion marker *Paxillin:*mKate (red) was injected in some embryos prior to cell dissociation. In control conditions, Yap-active cell colonies are surrounded by focal adhesions, while in Dasatinib-treated cells, focal adhesions fail to assemble around the colonies and they do no activate YAP (*TEAD:*Kaede in green). These images correspond to Supplementary videos 12-14. t=21h20min is shown in the images. Scale bars: 30 µm. **(D)** Still images from live imaging of dissociated zebrafish blastula cells from the reporter *TCF:*mCherry (red) cultured with DMSO (Control), the Wnt/β-catenin activator Chiron and the Wnt/β-catenin inhibitor IWR1. As expected, Chiron boosts *TCF*:mCherry activation intensity, while IWR1 inhibits it. These images correspond to Supplementary videos 15-17. t=26h40min is shown in the images. Scale bars: 30 µm. **(E)** Still images from live imaging of dissociated zebrafish blastula cells. Top panels show *TEAD:*Kaede activation in control vs. IWR1-treated cells; bottom panels display *TCF:*mCherry activity following Chiron activation alone or in combination with Dasatinib. This shows a reciprocal inhibition between these two signaling pathways. These images correspond to Supplementary videos 18-21. t=19h45min is shown in the top panel and t=16h48min in the bottom panel images. Scale bars: 30 µm. **(F)** In vivo effects of actomyosin contractility perturbations on YAP and β-catenin activation. Confocal images of optic cups from 24hpf zebrafish embryos expressing *TEAD:*Kaede (green, top) or *TCF:*mCherry (red, bottom) treated with DMSO, the ROCK inhibitor Rockout (to decrease tension), or Calyculin A (to hyperactivate actomyosin contractility). While decreased actomyosin contractility reduces both pathway signaling activation, its hyperactivation has no effect. Scale bars: 50 µm. **(G)** Still images from live imaging of dissociated zebrafish blastula cells from *TEAD:*Kaede embryos (green, top) treated with DMSO and Rockout or *TCF:*mCherry (red, bottom) treated with Chiron and a combination of Chiron and Rockout. The reduction of actomyosin tension also attenuates YAP and β-catenin signaling *in vitro*. These images correspond to Supplementary videos 22-25. t=19h45min is shown in the top panel and t=17h20min in the bottom panel images. Scale bars: 30 µm. **(H)** Still images from live imaging of dissociated zebrafish blastula cells from *TEAD:*Kaede embryos (green) and *TCF:*mCherry (red) treated with DMSO or Chiron and seeded in compliant 1% Matrigel or more rigid and more concentrated 20% Matrigel substrate. Panels show enhanced activation of control, Chiron-stimulated *TCF:*mCherry reporter and of control *TEAD:*Kaede, suggesting increased transcriptional response to stiffer ECM or to higher ligand concentrations. These images correspond to Supplementary videos 26-31. t=19h45min is shown in *TEAD:*Kaede and t=26h40min in *TCF:*mCherry images. Scale bars: 30 µm.

It is important to note that while proper eye morphology and kinetics were significantly altered, we never observed a complete arrest of the folding process (Supp Figure 1), which is in agreement with previous knowledge that, besides RPE cell flattening (Moreno-Mármol et al., 2021), there are complementary programs that altogether drive this eye folding event, such as basal constriction of the neural retinal cells (Nicolás-Pérez et al., 2016; Shidaye et al., 2016; 2017) and cell migration towards the future retinal domain during RIM involution (Heermann et al., 2015).

### Mechanosensitive regulation of β-catenin and YAP activity by tissue tension *in vitro* and in RPE cells *in vivo*

To determine whether the activation of β-catenin and YAP is directly controlled by tissue tension, we first validated the efficacy of our pharmacological inhibitors. Milder inhibition of YAP (Dasatinib) and β-catenin (IWR-1), which resulted in only a minor delay in optic cup folding, was sufficient to prevent the activation of the YAP/TEAD and β-catenin/TCF transgenic reporters (Figures 2A and 2B; Supp Videos 8-11). This result suggests that reporter activity is highly sensitive to signaling disruption, even before major optic cup folding defects occur. Consistent with this, we found that inhibition of these pathways did not impair RPE cell fate specification *per se*. The expression of canonical RPE markers, such as *bhlhe40* and *tfec*, remained unaffected, indicating that differentiation proceeded normally (Supp Figure 3A and 3B). Instead, the disruption specifically targets RPE geometry, suggesting that YAP and β-catenin signaling are required for the physical execution of morphogenesis rather than the establishment of tissue identity. To further isolate the signaling response from the morphogenetic formation of the eye, we established a primary culture of dissociated zebrafish blastula cells. When we treated these dissociated cells with the inhibitors Dasatinib or IWR-1, we observed a significant decrease of reporter activity, confirming that the disruption of these pathways is not a consequence of global eye morphogenesis impairments (Figure 2C and 2D; Supp videos 12-17). Notably, when visualizing Focal Adhesions (FAs) via *paxillin:mKate* expression, we observed that YAP-activating cell colonies were surrounded by a dense FA meshwork (Figure 2C; Supp video 13), which was completely abolished following Dasatinib treatment, consistent with its known role in blocking YAP activation by disrupting FA-mediated signaling (Figure 2C; Supp video 14). In the case of β-catenin/TCF reporter cells, we used a pharmacological activator, Chiron, to obtain higher and more reliable levels of activation intensity (Figure 2D; Supp video 16).

**Figure 3.**
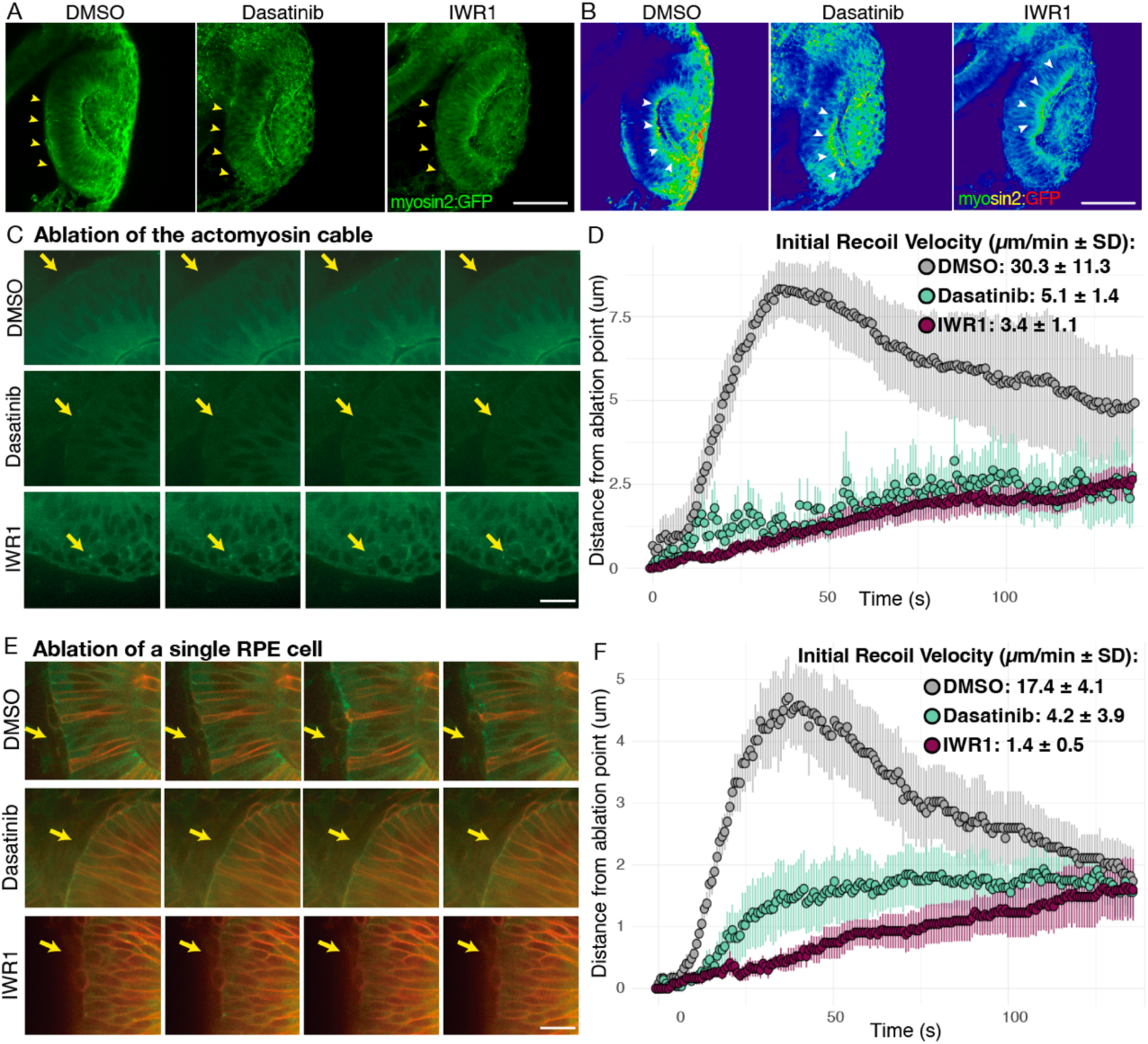
Inhibition of YAP and β-catenin reduces the relative tissue tension at the RPE-NR interface. **(A, B)** Confocal images (A) and corresponding heatmaps (B) of optic cups from 23hpf zebrafish *myosin2:*GFP reporter embryos treated with DMSO (control), Dasatinib, or IWR1. Yellow arrowheads in (A) and white arrowheads in (B) points at the actomyosin cable in the RPE-NR interface (A) and the NR-lens interface (B). Scale bars: 100 µm. **(C)** Time-lapse snapshots showing laser ablation of the actomyosin cable across DMSO, Dasatinib, and IWR1-treated embryos. Yellow arrows point to the cutting site. These images correspond to Supplementary videos 32-34. Scale bars: 30 µm. **(D)** Quantification of the distance (µm) from the ablation point over time (s), illustrating tissue relaxation. Text shows the initial recoil velocity (µm/min +/− SD) for every treatment. n=4 DMSO embryos; n=4 Dasatinib-treated embryos; n=7 IWR1-treated embryos. **(E)** Time-lapse snapshots showing laser ablation of a single RPE cell across DMSO, Dasatinib, and IWR1-treated embryos. Yellow arrows point to the cutting site. These images correspond to Supplementary videos 35-37. Scale bars: 30 µm. **(F)** Quantification of the distance (µm) from the ablation point over time (s), illustrating tissue relaxation. Text shows the initial recoil velocity (µm/min +/− SD) for every treatment. n=5 DMSO embryos; n=4 Dasatinib-treated embryos; n=6 IWR1-treated embryos.

Using cultured dissociated blastula cells, we observed that treatment with either Dasatinib (YAP inhibitor) or IWR-1 (β-catenin inhibitor) resulted in the cross-inhibition of the opposing pathway. This means that inhibiting YAP reduced β-catenin reporter activity, while the inhibition of β-catenin similarly decreased YAP activation (Figure 2E; Supp videos 18-21). This reciprocal inhibition strongly suggests the existence of a robust positive crosstalk regulation loop between these two pathways in the RPE during eye development. This reciprocal activation between both pathways is consistent with previous reports in other systems showing that YAP/TAZ are integral components of the β-catenin destruction complex (Azolin et al., 2014) and that β-catenin can directly regulate YAP expression (Heallen et al., 2011; Rosenbluh et al., 2012).

Next, we manipulated actomyosin-mediated intracellular tension *in vivo* using Rockout (a contractility inhibitor) and Calyculin A (a contractility enhancer). Consistent with our previous observations, activation of both the YAP and β-catenin pathways was prevented exclusively by reducing actomyosin tension (Figure 2F). In contrast, enhancing contractility with Calyculin A did not significantly alter the signaling levels (Figure 2F), likely because the RPE cellular morphology and optic cup folding were already at a mechanical maximum under control conditions (Figures 1D, 1F, and 2F). To decouple pathway activation from the global morphogenetic process, we performed cell culture experiments and treated dissociated YAP and β-catenin sensor cells with Rockout. We observed that signaling activation was intrinsically sensitive to actomyosin tension, even independently of the defect in eye morphogenesis *in vivo* (Figure 2G, Supp videos 22-25). Finally, to manipulate the cellular mechanical properties via an independent strategy, we increased the concentration of the Matrigel substrate coating twenty-fold (20X). This higher concentration led to a robust increase in the number and size of cellular colonies showing YAP and β-catenin activation (Figure 2H; Supp videos 26-31). It is important to note that while this suggests mechanosensitive induction, this effect may result from an increase in substrate stiffness, ligand density, or a combination of both (Anguiano *et al*, 2020; Wu *et al*, 2025).

### YAP and β-catenin are required to maintain actomyosin-mediated tissue tension at the RPE-retinal interface

To investigate whether YAP and β-catenin signaling not only respond to changes in tissue tension but also actively maintain it, we evaluated actomyosin-dependent tension in the eye following pathway inhibition. In the zebrafish optic cup, two distinct high-tension actomyosin networks are essential for folding: one at the basal surface of the neural retina (NR) and the other at the apical interface between the NR and RPE (Nicolás-Pérez *et al*, 2016; Sidhaye & Norden, 2017; Moreno-Mármol *et al*, 2021).

Using the *Myl12.1:eGFP* reporter line (Behrndt *et al*, 2012), we observed that while the basal NR actomyosin cable remained largely intact after treatment with Dasatinib and IWR-1, the apical cable at the RPE-NR interface was severely attenuated (Figure 3A-B). This highlights the tissue-specific requirement for these pathways within the RPE, where they are exclusively active (Figure 1A, 1B). This localized reduction in actomyosin intensity suggests a corresponding decrease in tissue tension.

To quantify this, we performed laser ablation experiments to measure the instantaneous recoil velocity of RPE (Zulueta-Coarasa & Fernandez-Gonzalez, 2017; Vogel & Venugopalan, 2003). We targeted two locations: the apical actomyosin cable at the RPE-NR interface and a single RPE cell body. In control (DMSO-treated) animals, we observed a considerably high initial recoil velocity at both locations (Figure 3D, 3F; Supp videos 32, 35). This corroborates that the RPE is maintained in a state of high endogenous tension during morphogenesis, with the actomyosin cytoskeleton actively generating the forces required for the expansion and bending of the optic cup. This high mechanical baseline is consistent with the stretching and basal constriction reported during RPE development (Moreno-Mármol *et al*, 2021).

Following the inhibition of YAP and β-catenin, the recoil velocity was dramatically reduced, suggesting that the tissue loses mechanical tension (Figure 3D, 3F; Videos 33, 34, 36, 37). To ensure that the observed reduction in RPE tissue tension was a direct consequence of impaired mechanotransduction rather than altered tissue growth or viability, we assessed the proliferative and apoptotic profiles of RPE cells after inhibition of the YAP and β-catenin pathways. We used phospho-Histone3 (mitotic marker) and Caspase 3 (apoptotic marker) antibodies and confirmed that, as in control RPE cells (DMSO), pharmacologically interfered embryos showed highly stable RPE tissue with little cell proliferation and almost negligible cell death (Supp Figure 3C-3F). Furthermore, the maintenance of canonical RPE markers in the inhibition experiments (Supp Figure 3A and 3B) and the relative, although delayed, completion of optic cup folding (Supp Figure 1) suggest that the RPE remains a viable epithelial sheet, specifically impaired in its mechanical ability to generate or respond to tissue-level tension (Figure 2 and Figure 3). Our observations are consistent with previous results showing that RPE expansion during these stages relies on coordinated cellular stretching rather than changes in cellular density (Hernández-Bejarano *et al*, 2015; Moreno-Mármol *et al*, 2021), and support the notion that the decrease in recoil velocity in the RPE of YAP- and Wnt-perturbed embryos is related to a decrease in actomyosin-generated stress. Therefore, we can conclude that YAP and β-catenin function within a self-sustaining mechanical loop in the RPE: they sense environmental tension and subsequently signal the cell to maintain the actomyosin contractility required for mechanical homeostasis and successful optic cup folding.

### Transcriptomic profiling reveals a shared gene regulatory program between YAP and β-catenin pathways governing the RPE mechanical landscape

To identify the molecular targets downstream of the YAP and β-catenin mechanical loop, we performed comparative transcriptomic profiling of embryos following pharmacological inhibition of YAP and β-catenin. Kyoto Encyclopedia of Genes and Genomes (KEGG) enrichment analysis of the Differential Gene Expression in embryos treated with Dasatinib (YAP inhibitor), IWR-1 (β-catenin inhibitor), Rockout (actomyosin contractility inhibitor) and Yoda1 (dysregulated mechanosensitivity), showed a significant clustering around cytoskeletal dynamics, cell-cell junctions, and cell-ECM adhesions (in red; Figure 4A), as well as common signaling pathways, such as the MAPK and Wnt (in yellow; Figure 4A) and in known mechanosensitive processes, such as Calcium signaling pathway and Endocytosis (in blue; Figure 4A). Interestingly, the KEGG of Endocytosis was found to be highly significant in all the inhibitory treatments (Figure 4A). Endocytosis is a mechanosensitive process that balances membrane tension and the surface area. The downregulation of the endocytic machinery, together with the loss of actomyosin contractility (Figure 3) after inhibiting these pathways, suggests that the YAP and β-catenin circuits might allow the RPE to accommodate the rapid increase in surface area required for optic cup folding without compromising epithelial integrity. Consistent with KEGG enrichment analysis, we also performed a Gene Ontology (GO) enrichment analysis of Cellular Components (CC), and found a highly significant enrichment of terms associated to anchoring junctions and adherens junctions across all experimental conditions (Supp Figure 4A-D).

**Figure 4.**
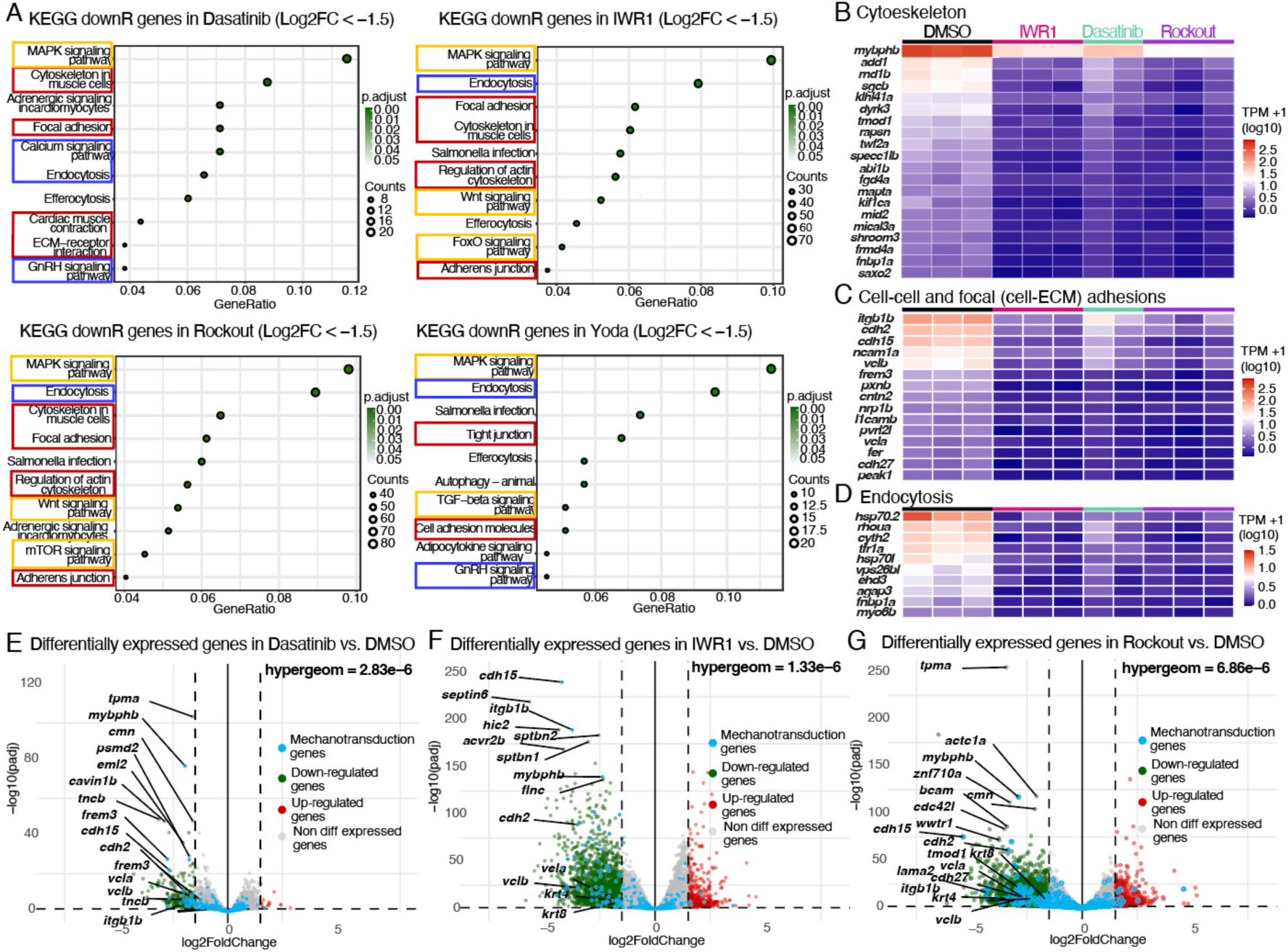
Transcriptomic analysis shows reduction of mechanostransduction-related genes following reduction of intracellular tension and YAP/β-catenin signaling. **(A)** KEGG pathway enrichment plots of down-regulated genes (with Log_2_ FoldChange ≤ −1.5) in embryos treated with Dasatinib, IWR1, Rockout, or Yoda compared to DMSO controls. The x-axis indicates gene ratio, dot size represents the gene count within each pathway, and dot color indicates the adjusted p-value. Key structural cytoskeletal and adhesion (red), mechanosensitive (blue) and signaling pathways (yellow) are highlighted by colored boxes. **(B-D)** Heatmaps displaying the expression levels (TPM+1 log_10_ scale) of representative common down-regulated genes between IWR1, Dasatinib, and Rockout-treated groups compared to the control DMSO group. Genes are grouped by functional annotations: Cytoskeleton **(B)**, Cell-cell and focal (cell-ECM) adhesions **(C)**, and Endocytosis **(D)**. **(E-G)** Volcano plots showing differentially expressed genes (DEGs) in Dasatinib vs. DMSO **(E)**, IWR1 vs. DMSO **(F)**, and Rockout vs. DMSO **(G)**. y-axis represents the negative log_10_ of the adjusted p-values and x-axis represents log_2_ FoldChange. Down-regulated genes (green), up-regulated genes (red), non-differentially expressed genes (grey), and specific mechanotransduction-related genes (blue) are shown. Statistical significance for the enrichment of mechanotransduction genes is indicated by the hypergeometric test (top right of each plot). Key down-regulated genes are labeled.

To confirm the specificity of the different pharmacological inhibitors, we performed a hypergeometric test to determine whether the downstream targets were statistically downregulated after the different treatments (Falcon & Gentleman, 2007) (Supp Figure 4E-G). We observed a statistically significant downregulation of pathway-specific targets in Dasatinib, IWR1 and Rockout treatments. However, while Yoda 1 is widely used as an upstream regulator of mechanosensation via Piezo 1, we were unable to validate the specificity of this treatment in our biological context. This was evidenced by the lack of statistically significant differential expression among downstream ion channel-related genes following Yoda1 treatment. For this reason, Yoda 1 treatment was excluded from subsequent analysis.

To specifically isolate the mechanisms governing RPE mechanical homeostasis, we focused on the convergent targets of actomyosin contractility and the YAP/β-catenin pathways. Transcriptomic screening revealed that the KEGG pathways obtained following YAP and β-catenin inhibition closely mirrored those obtained following direct actomyosin disruption via Rockout. We conducted a targeted analysis of the most significant differentially expressed genes (DEGs) shared across the Dasatinib, IWR-1, and Rockout datasets (Figures 4B-4D). Within the cytoskeleton-related KEGG pathways, we identified the downregulation of genes essential for actomyosin contractility and stability, including myosin-binding proteins (*mybphb*), actin-binding regulators (*tmod*, *twf2a*), and microtubule-associated proteins (*mapta*, *mical3a*, *kif1ca*). Furthermore, the downregulation of Rho-family GTPase regulators, such as *rnd1b* and *fgd4*, indicates a collapse in the upstream signaling required for cytoskeletal tension (Figure 4B). Consistent with the observed loss of tissue integrity, we found a significant decrease in the transcripts for cell-cell and focal adhesions. This included multiple E-cadherin isoforms (*cdh2*, *cdh15*, *cdh27*) and contactin (*cntn2*), along with core focal adhesion components such as integrin beta 1 (*itgb1*), vinculins (*vcla*, *vclb*), and paxillin (*pxnb*) (Figure 4C). The coordinated downregulation of these structural anchors likely explains the severe reduction in mechanical recoil and attenuation of the apical actomyosin cable observed following pathway inhibition (Figure 3). Finally, we also found common downregulated genes related to the endocytosis mediated membrane trafficking, such as the Ras homolog family member *rhoua*, formin binding protein *fnbp1a* and regulators of GTPase-activator proteins *cytohesin-2* and *agap3* (Figure 4D). Interestingly, this subset of genes identified within the KEGG pathways associated with the cytoskeleton, cell adhesions, and endocytosis (termed mechanotransduction genes), were predominantly downregulated upon pathway inhibition, with only a marginal or nonexistent number of genes being upregulated (Figures 4E-4G). Given the higher total number of downregulated genes across the dataset (Supp Figure 4H), we performed a hypergeometric test to determine whether this enrichment was statistically significant (Falcon & Gentleman, 2007). For all three treatments (Dasatinib, IWR-1, and Rockout), the highly significant p-values confirmed that the downregulation of these mechanotransduction genes was a targeted effect of the inhibitors and not a mere consequence of the overall gene distribution (Figures 4E-4G). Interestingly, in contrast to the group of downregulated genes, the subset of common upregulated genes across all three treatments did not reveal any enriched KEGG pathway associated to mechanotransduction (Supp Figure 4H-J).

By combining the physical measurements of tissue tension with statistically rigorous transcriptomic profiling, we demonstrated that YAP and β-catenin are not merely sensors of the mechanical environment but are essential transcriptional drivers of the architecture of the RPE. By co-regulating the expression of the actomyosin engine, its membrane anchors, and elasticity (endocytosis machinery), this mechanochemical loop ensures that the RPE maintains the homeostatic tension and adhesive strength necessary to drive optic cup folding.

### The LINC complex might act as the mechanotransducer between RPE stretching and YAP and β-catenin activation

Finally, we sought to identify the molecular transducer that might convert cellular stretching into nuclear activation of YAP and β-catenin, specifically within the RPE. We focused on the Linker of Nucleoskeleton and Cytoskeleton (LINC) complex, a conserved protein bridge known to transmit mechanical forces from the cytoplasmic actomyosin network directly to the nuclear envelope to regulate gene expression. The LINC complex is composed of large KASH-domain proteins, the Nesprins, located in the outer nuclear membrane, where they bind directly to cytoplasmic actin filaments, microtubules, and motor proteins. The KASH domain of Nesprins hooks onto SUN proteins in the inner nuclear membrane, which act as anchors for the nuclear lamina and chromatin-binding proteins (Khilan *et al*, 2021). We initially performed CRISPR-mediated knockout (KO) of three individual zebrafish Nesprin paralogs (*syne1a*, *syne1b*, and *syne3*). The initial screening at F0 did not show any correlation between the mutation of these three Nesprin paralogs and the overt phenotype of the fish embryos. We also generated stable KO lines for *syne1b* and *syne3*, but they also failed to produce a phenotype, likely due to genetic compensation or redundancy, as previously reported in mouse *Nesprins1/2* mutants (Zhang *et al*, 2007). To circumvent this, we shifted our strategy to the expression of dominant-negative (dn) forms of the three Nesprin paralogs (*nes* DN) <u>(Zhang *et al,* 2001; Elosegui-Artola *et al*, 2017)</u>. Global embryo expression causes severe developmental arrest before optic cup formation (Supp Figure 5A, 5B). Despite attempts to titrate the dose, we could not find a concentration that avoided this global arrest in the overall development. Consequently, we designed targeted transplantation experiments to inhibit Nesprin expression more specifically in the eye progenitor population. We injected dominant-negative (DN) forms of the three nesprins (*nes* DN) into a donor embryo and transplanted the uppermost animal pole blastoderm cells (highly enriched in eye progenitor cells) into a WT host embryo (Figure 5A, 5F, and 5K). Our first observation was that when Nesprin-inhibited cells populated the RPE domain (green cells with a red nuclear carrier in Figures 5B-5E), they failed to undergo the characteristic stretching and flattening required for morphogenesis (Figures 5D, 5E). Crucially, this effect was tissue-specific; when Nesprin function was inhibited exclusively in the Neural Retina (NR), eye folding proceeded normally (Figure 5B). We observed a strikingly similar phenotype when transplanting progenitor cells expressing a *yap* DN form (cells with a red nuclear carrier in Figures 5G-5J), where RPE-specific inhibition of YAP prevented cellular flattening and subsequently altered eye folding (Figure 5J). Additionally, a high population of YAP-inhibited cells in the RIM region appeared to interfere with proper RIM involution (Figure 5I). Finally, we used YAP and β-catenin activity reporter lines to determine whether the LINC complex is required for nuclear activation of these pathways. By transplanting *nes* DN cells into reporter hosts (red cells in the YAP/TEAD:GFP reporter line and green cells in the β-catenin /TCF:mCherry reporter line), we confirmed that RPE cells lacking functional Nesprins failed to activate either YAP or β-catenin signaling (Figures 5L-5O). These results suggest that the LINC complex is an indispensable physical bridge that translates the high mechanical strain of the RPE into the activation of the RPE morphogenetic and transcriptional program.

**Figure 5.**
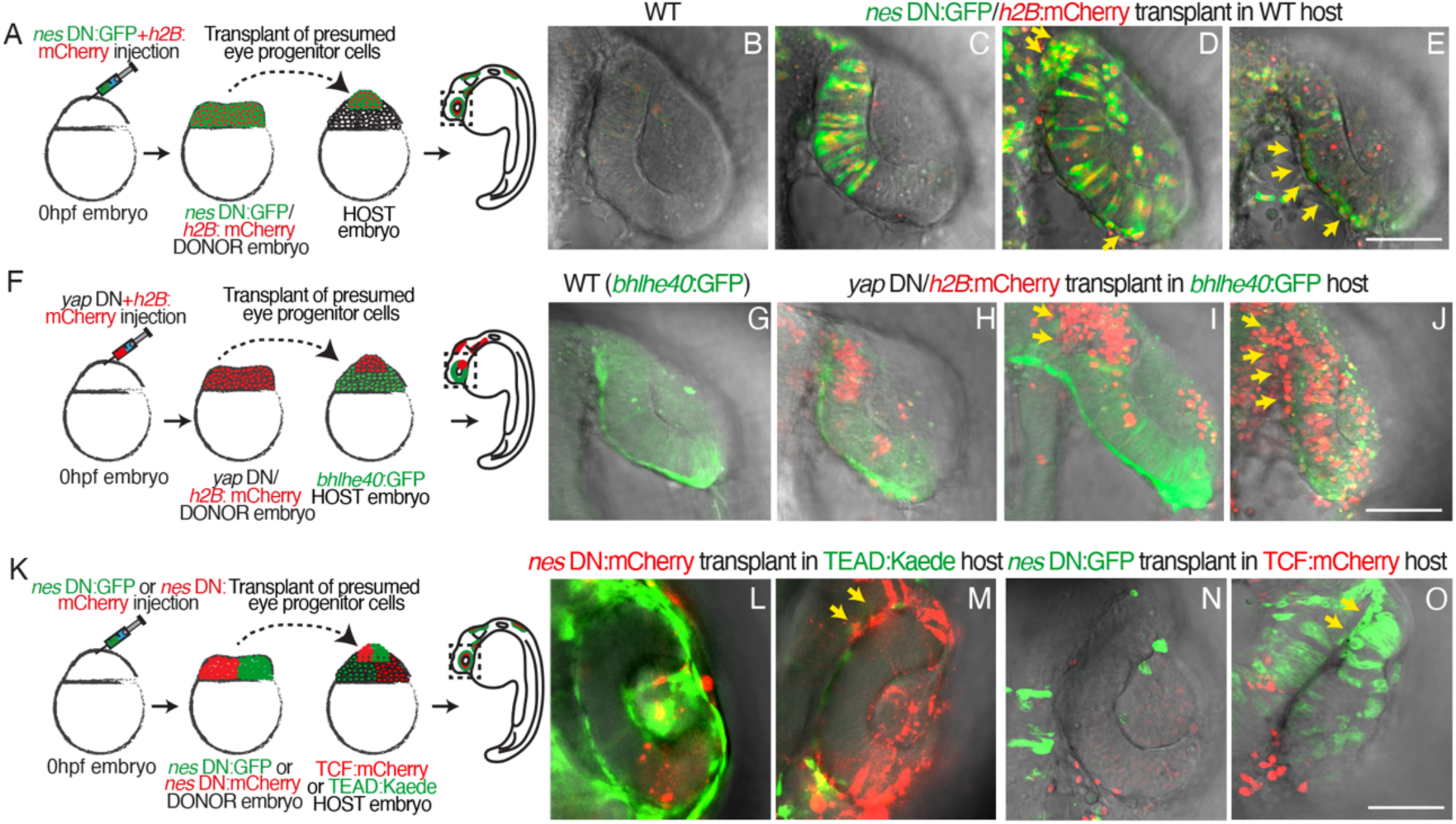
Mosaic inhibition of the LINC domain reveals a specific effect and requirement for YAP/β-catenin activation in RPE cells. **(A-E)** Transplantation of Nesprin dominant-negative (*nes* DN) cells from a donor into WT host. **(A)** Schematic outline of the experiment. Donor embryos are injected at 0hpf stage with *nes* DN:GFP and *h2B:*mCherry mRNAs, and presumed eye progenitor cells (from the most animal region) are subsequently transplanted into WT host embryos at sphere stage. Confocal images of a control WT optic cup **(B)** and of *nes* DN:GFP/*h2B:*mCherry donor cells (green cells with a red nuclear carrier) integrated into a WT host optic cup **(C-E)**. No effect on optic cup folding and cellular geometry was seen when nesprin-inhibited cells populated just the NR domain **(C)**. In contrast, when nesprin-inhibited cells populate the RPE, flattened cellular geometry **(D)** and optic cup folding are affected **(E)**. Yellow arrows highlight nesprin-inhibited RPE cells with altered geometries. Scale bars: 50 µm. **(F-J)** Transplantation of YAP dominant-negative (*yap* DN) cells from a donor into *bhlhe40*:GFP reporter WT host (Moreno-Mármol *et al*, 2021). **(F)** Schematic outline of the experiment. Donor embryos are injected at 0hpf stage with *yap* DN and *h2B:mCherry* mRNAs, and presumed eye progenitor cells (from the most animal region) are subsequently transplanted into WT *bhlhe40*:GFP reporter host embryos at sphere stage. Confocal images of a control *bhlhe40*:GFP reporter WT optic cup (RPE domain in green) **(G)** and of *yap* DN/*h2B:*mCherry donor cells (red cells) integrated into a WT *bhlhe40*:GFP reporter host optic cup **(H-J)**. No effect on optic cup folding and cellular geometry was seen when yap-inhibited cells populated just the NR domain **(H)**. In contrast, when nesprin-inhibited cells populate the RPE, cells will accumulate at the RIM region **(I)** and optic cup folding is affected **(J)**. Yellow arrows highlight yap-inhibited RPE cells with altered geometries. Scale bars: 50 µm. **(K-O)** Effects of *nes* DN donor clones on YAP and β-catenin signaling. **(K)** Schematic outline of the experiment. Donor embryos are injected at 0hpf stage with either *nes* DN:mCherry or *nes* DN:GFP mRNAs and presumed eye progenitor cells (from the most animal region) are subsequently transplanted into WT *TEAD:*Kaede **(L,M)** or *TCF:*mCherry **(N,O)** reporter host embryos at sphere stage. Confocal images of a control WT *TEAD:*Kaede (green) **(L)** and *TCF:*mCherry (red) reporter optic cup (N). (M) *nes* DN:mCherry donor cells (red) transplanted into a *TEAD:*Kaede reporter host (green). Clones of nesprin-inhibited cells failed to activate YAP pathway. (O) *nes* DN:GFP donor cells (green) transplanted into a *TCF:*mCherry reporter host (red). Clones of nesprin-inhibited cells failed to activate β-catenin pathway. Yellow arrows point to the nesprin-inhibited clones of cells. Scale bars: 50 µm.

## Discussion

### YAP and β-catenin signaling is specifically activated in the RPE cells probably due to their flattened geometry and the LINC complex

Our results demonstrate that the YAP and β-catenin pathways are specifically activated within the RPE layer of cells, but remain absent in adjacent eye populations such as the neural retina (NR). This is evidenced by the restricted activation of YAP and β-catenin reporters in the RPE (Figure 1A, 1B), alongside by the enrichment of downstream targets and effectors of both β-catenin and YAP signaling in RPE clusters relative to the NR (Figure 1C). Beyond differing in tissue identity, the RPE and NR display distinct mechanical and morphogenetic profiles: the RPE features a significantly higher degree of rigidity (Buono *et al*, 2021; Moreno-Mármol *et al*, 2021) and adopts a markedly elongated, flattened geometry, contrasting with the cuboidal shape of NR cells (Sidhaye & Norden, 2017).

Intriguingly, our findings link the distinct rigidity, intracellular tension, and geometry of RPE cells to the specific activation of these YAP and β-catenin cascades. Supporting the connection to cellular tension, we made several key observations: (i) the activation of both pathways is prevented when actomyosin contractility is pharmacologically reduced using Rockout (Figure 2F); (ii) inhibiting these pathways severely attenuates the high-tension actomyosin cable specifically at the apical RPE-NR interface, while leaving the NR-lens basal cable intact (Figure 3A, 3B); and (iii) when signaling is decoupled from global eye development using primary zebrafish blastula cell culture, pathway activation remains intrinsically sensitive to intracellular tension (Figure 2G) and probably also substrate stiffness (Figure 2H).

Finally, supporting a direct link between pathway activation and cellular geometry, we explored the involvement of the LINC complex. Mosaic transplantation of cells expressing dominant-negative forms of Nesprins (*nes* DN) revealed that RPE cells are uniquely sensitive to LINC-mediated mechanotransduction. While *nes* DN-expressing RPE cells failed to undergo characteristic stretching, *nes* DN-expressing NR cells maintained their typical cuboidal shape without hindering optic cup morphogenesis (Figure 5B–E). Crucially, these transplanted, LINC-deficient RPE cells also failed to activate either the YAP or β-catenin signaling reporters.

These observations strongly suggest that the physical strain of RPE cellular stretching is translated via the LINC complex to trigger the nuclear translocation and activation of these transcription factors. Mechanistically, this tissue specificity aligns with the distinct physical forces driving eye morphogenesis; while the NR drives folding primarily through localized basal constriction (generating pushing and bending forces), the RPE undergoes massive lateral stretching (generating pulling forces). The LINC complex is uniquely suited to sense this lateral strain because flattening a cell physically deforms the nucleus, altering nuclear pore mechanics to force transcription factors inside (Elosegui-Artola *et al*, 2017; Thorpe & Lee, 2017; Koushki *et al*, 2023).

In summary, our results on the involvement of the LINC complex in the activation of YAP and β-catenin specifically in the elongated RPE cells help to close the loop, where the LINC complex senses high basal tissue tension to contribute to nuclear activation of the YAP and β-catenin transcription factors. There, they will drive the expression of adhesion and actomyosin genes to maintain the cellular rigidity and the actomyosin cable in the RPE/NR interface, which is key to maintain and promote the RPE cellular stretching to contribute to the folding of the eye.

### Reciprocal YAP and β-catenin crosstalk drive the homeostatic tension required for optic cup folding

A striking finding of our study is the profound functional convergence and transcriptional overlap between the YAP and β-catenin pathways during RPE morphogenesis, despite their distinct molecular upstream and downstream pathways of activation. We have observed several differences in their activation, such as that while β-catenin follows a transient wave of activation (Figure 1A), probably driven by localized *wnt2* expression (Supp Figure 2), YAP displays a sustained activation pattern (Figure 1B), which might be coupled to focal adhesion assembly (Figure 2C). However, the genetic or pharmacological disruption of either pathway leads to a very similar morphogenetic failure, which is the reduction of the apical actomyosin cable at the RPE-NR interface and impaired RPE cell flattening (Figure 1D). In the same way, the inhibition of either pathway leads to a significant downregulation of genes that co-regulate a similar mechanotransduction gene program (Figure 4). This might be explained by the reciprocal positive feedback loop that is backed up by our cell culture experiments, where we can see that the inhibition of one pathway seems to abolish the activation of the other (Figure 2E). Therefore, besides their specificities, rather than acting as redundant signaling cascades, YAP and β-catenin appear to operate within a self-sustaining co-regulatory circuit.

In this circuit, LINC-mediated physical stretch of the nuclear pores may initiate the nuclear translocation of both factors, which then cooperatively maintain each other into an active state. This reciprocal stabilization ensures a unified and amplified transcriptional response, driving the regulation of the adhesion and cytoskeletal components required to sustain high tissue-level tension in the RPE during optic cup folding. According to this, it has been shown that YAP/TAZ are components of the β-catenin destruction complex under low cortical tension or the absence of Wnt activating ligands. Conversely, when tension increases, the β-catenin destruction complex is disrupted and both transcription factors (YAP and β-catenin) are free to enter the nucleus together (Azzolin *et al*, 2014). Moreover, it has been shown that β-catenin can directly bind to the promoter of *yap1* to bust its expression (Rosenbluh *et al*, 2012) and that active nuclear YAP/TEAD complexes bind to the enhancers and promoters of *wnt* ligands and *frizzled* receptors, boosting the sensitivity of the cell to Wnt signaling (Park *et al*, 2015; Gregorieff *et al*, 2015). Finally, it has been shown that YAP can enter though the nuclear pores via the LINC complex that transmits the actomyosin tension to the nucleus (Elosegui-Artola *et al*, 2017).

### The mechanical gene regulatory program of YAP and β-catenin maintains RPE morphogenesis and optic cup folding

Our transcriptomic profiling following pharmacological inhibition support the existence of this sustained transcriptional program activated by YAP and β-catenin pathways to actively maintain high-basal intracellular tension in the RPE cells, where these pathways are specifically active. Our differential gene expression, KEGG pathway and GO enrichment analyses revealed high convergence among RNAseq datasets derived from embryos treated with Dasatinib (YAP inhibitor), IWR-1 (β-catenin inhibitor), and Rockout (actomyosin contractility inhibitor). The finding that downstream transcriptional profiles following YAP and β-catenin inhibition closely mirror the effects of direct intracellular tension disruption implies that these pathways might operate as core components of an integrated circuit, functioning as essential transcriptional drivers that build and maintain the mechanical machinery of the RPE cells. A detailed analysis of the shared DEGs uncovers an interference of the structural networks required to generate and sustain tissue-level tension. Within the cytoskeleton-related KEGG pathways, we found that YAP, β-catenin and intracellular tension inhibition led to a significant downregulation of factors crucial for actomyosin contractility, structural stability and its regulators. This includes the downregulation of myosin-binding proteins (e. g. *mybphb*), actin-binding regulators (e. g. *tmod* and *twf2a*), microtubule-associated components (e. g. *mapta, mical3a* and *kif1ca*) and of Rho-family GTPase regulators (e. g. *rnd1b* and *fgd4*) as upstream signaling driving actomyosin network failure. Importantly, this transcriptional program controlled by YAP and β-catenin signaling might explain the “softening” of the RPE cells and the dramatic reduction in mechanical recoil velocity of the actomyosin cable at the RPE-NR interface observed after inhibiting these signaling pathways (Figure 3).

This transcriptional control goes beyond the intracellular actomyosin network to the physical anchors that transmit forces across cell-to-cell boundaries and cell-to-ECM. We also observed a reduction in transcripts encoding cell-cell and focal (cell-ECM) adhesion components. This includes multiple E-cadherin (e.g. *cdh2, cdh15* and *cdh27*), contactin (*cntn2*), and other core components of the focal adhesions, such as integrin (e.g. *itgb1*), vinculins (e.g. *vcla* and *vclb*), and paxillin (e.g. *pxnb*). Statistical analysis using hypergeometric tests confirmed that this systematic downregulation of mechanotransduction-related genes was specific, with virtually no mechanical or structural genes present in the common upregulated gene datasets. We hypothesize that without the molecular anchors, the RPE would lose its ability to distribute and balance physical forces across the epithelial sheet. Thus, the YAP and β-catenin self-sustaining program acts as a transcriptional master regulator, ensuring that the cellular actomyosin engine and its adhesions are co-regulated to robustly guide RPE tissue properties and optic cup morphogenesis. It is also important to note that after inhibiting YAP and β-catenin pathways, the vast majority of the genes are downregulated (90% in Dasatinib treatment, 80% in IWR1 treatment), which might indicate that these signaling pathways are positive transcriptional activators of the structural mechanical engine of the RPE cells, and not general modulators of RPE and/or eye development. Consistent with this, we do not find enrichment of any specific pathway among the upregulated gene datasets (Supp Figure 4I), neither general KEGG terms such as “eye development” or “eye morphogenesis”, strengthening the argument that YAP and β-catenin pathways are specifically responding to high tensional states by maintaining the mechanical engine of the RPE cells, rather than regulating general identity programs.

Finally, our transcriptomic analysis also uncovers an interesting and fairly unexplored dimension to this coordinated mechanical loop, which is the highly significant downregulation of components of the endocytic machinery across all treatments (YAP and β-catenin inhibition, piezo-related mechanosensitivity and the reduction of actomyosin contractility) (Figure 4D). While endocytosis is traditionally viewed as a mechanism of cellular trafficking, it also entails a mechanosensitive process required to balance membrane tension and surface area. RPE cells transition from rounded geometries into highly stretched and flattened configurations, while they experience strong lateral strain and a considerable expansion of surface area. The downregulation of endocytosis genes upon the inhibition of YAP and β-catenin, suggests that these pathways might increase endocytic rates to act as a mechanical buffering system that allows the membranes to accommodate the dramatic geometrical changes that the RPE experience without compromising structural and epithelial integrity.

### YAP and β-catenin act as a mechanical rheostat to maintain RPE cellular contractility

Our findings suggest that YAP and β-catenin operate as transcriptional co-activators of a highly dynamic mechanical rheostat within the developing RPE. During eye morphogenesis, complex tissue remodeling and cell movements introduce fluctuating mechanical strains across the epithelial sheet. Rather than acting as simple on/off molecular switches, RPE-restricted YAP and β-catenin activity continuously monitors and adjusts actomyosin cortical tension and cellular adhesions. This sensitivity allows the cells to adapt to both extrinsic forces from the surrounding tissues and intrinsic forces generated within the RPE itself. By maintaining this continuous mechanochemical feedback loop (Hannezo & Heisenberg, 2019; De Santis *et al*, 2025), the tissue ensures that RPE cells achieve and hold the precise degree of elongated, flattened geometry required to physically drive optic cup invagination.

Several lines of evidence support this mechanical rheostat model based on YAP/β-catenin sustained mechanical feedback loops, defining both its baseline operating conditions and its dynamic range. First, laser ablation of the apical actomyosin cable and the single RPE cell body in control embryos reveals a considerably high initial recoil velocity (Figure 3C-3F). This inferred physical measurement confirms that the tissue operates under strong endogenous tension. This high basal mechanical tension defines the specific “operational range” of the rheostat, showing that the system is adapted to function within a high-force regime. Reflecting an upper limit or “maximum capacity” setting of this rheostat, amplifying actomyosin contractility via Calyculin A treatment yields no further change in cellular geometry or folding kinetics, suggesting that the system normally operates at its mechanical peak (Figure 1D and Figure 2F). Second, our data demonstrates that this rheostat possesses a wide dynamic range of action, meaning that YAP and β-catenin signaling are not statically fixed but are responsive to a broad spectrum of physical inputs. The pathways should be able to sense cortical tension of the RPE cells to activate just as the cells undergo pronounced elongation. Furthermore, the YAP/β-catenin rheostat responds to lowering cortical tension via Rockout treatment (Figure 2F), and also to changes in the concentration of the extracellular matrix surfaces (Figure 2H). Third, by functioning as a rheostat that tunes the physical properties of the tissue, these pathways act as speed regulators for morphogenesis. When we pharmacologically or genetically disrupt this fine-tuning, the rate of development is altered, explaining the consistent delay in optic cup folding kinetics. Crucially, because there are complementary morphogenetic programs at play, this disruption delays rather than completely arrests the process (Figure S1), proving that the rheostat specifically dictates mechanical efficiency. Finally, we have shown that YAP and β-catenin are linked in a reciprocal feedback loop, stabilizing the entire rheostat by acting as a molecular buffer that prevents stochastic fluctuations in signaling from causing structural failures during eye formation (Figure 2E). In summary, the concept of a mechanical rheostat (Weng *et al*, 2016; Vining & Mooney, 2017) conceptualizes how YAP and β-catenin interpret the high-tension state of developing RPE cells, turning physical strain into a sustained transcriptional program that actively maintains and fine-tunes the morphogenetic forces shaping the optic cup.

## Materials and Methods

### Zebrafish husbandry and mRNA microinjection

Embryos from the transgenic line *Tg(b-actin:myo:GFP)* were used to visualize the actomyosin network. To label the cell membranes and identify retinal pigment epithelium (RPE) cells, embryos at the 1-cell stage were microinjected with1nL of *Lyn-dTomato* mRNA at a concentration of 50 ng/µL. Following injection, embryos were incubated at 28.5°C in E3 medium supplemented with 0.0001% methylene blue to prevent fungal growth.

Zebrafish wild-type strains and the transgenic lines Tg(4X*GTIIC:*Kaede) (kindly provided by Pujades Lab at UPF), Tg(7x*TCFsiam:*mCherry) (kindly provided by Moro Lab at UNIPD), Tg(E1-*bhlhe40*:GFP) (kindly provided by Bovolenta Lab at CBMSO) and *Tg(b-actin:myl12.1:*eGFP) (kindly provided by Heisenberg Lab at ISTA) were maintained under standard husbandry conditions. All animal experiments were conducted according to institutional ethical guidelines and approved by the Animal Experimentation Ethics Committees of University Pablo de Olavide under license number 02/04/2018/041.

Embryo microinjections were performed at one-cell stage using a M400 Narishige microinjector with the injection volume calibrated using a calibration device. mRNAs for zebrafish *nes1a* DN:GFP, *nes1b* DN:GFP*, nes3* DN:GFP were injected together (*nes* DN:GFP) (100-150pg of each *nesprin* paralog in Supplementary Figure 5). mRNAs for zebrafish *nes1a* DN:mCherry, *nes1b* DN:mCherry*, nes3* DN:mCherry were injected together (*nes* DN:mCherry) (100-150pg of each *nesprin* paralog in Supplementary Figure 5). mRNA for *yap* DN was injected at 100-150 pg/embryo in Supplementary Figure 2. For the transplant experiments in Figure 5, 300 pg/embryo of mRNAs for *nes1a* DN:GFP, *nes1b* DN:GFP*, nes3* DN:GFP, *nes1a* DN:mCherry, *nes1b* DN:mCherry*, nes3* DN:mCherry and *yap* DN were injected in the donor embryos. mRNAs for *H2b*:mCherry, *H2b*:GFP, *utrophin:*GFP (Nicolás-Pérez *et al*, 2016), paxillin:mKate (Sidhaye & Norden, 2017) and *Lyn-dTomato* were injected at 50 pg/embryo. Following injection, embryos were incubated at 28.5°C (where necessary to slow developmental speed, embryos were cultured at 25°C) in E3 medium supplemented with 0.0001% methylene blue to prevent fungal growth.

For mRNA synthesis, DNA plasmids were linearized using NotI and then transcribed using the mMESSAGE mMACHINE SP6 Kit (AM1340, Thermo Fisher Scientifc) to synthesize capped mRNA. Injected embryos were mounted for in vivo imaging, dissociated for cell culture oz zebrafish blastula cells or fixed in 4% Formaldehyde (FA) along with their respective controls at the appropriate developmental stage.

Zebrafish constructs for *nesprins* and *yap* DN were designed following (Zhang *et al*, 2007).

### Dechorionation and Pharmacological Treatments

At 8 hours post-fertilization (hpf), embryos were enzymatically dechorionated using a solution of 30 ng/µL pronase in E3 medium for 10 minutes. Following thorough washing to remove the pronase, embryos were arrayed into plates (50 embryos per plate) and treated with the specified pharmacological inhibitors: IWR-1 (25 µM), Dasatinib (100 µM), Rockout (150 µM), Calyculin A (0.5 µM) and Yoda 1 (10 µM). In Figure 2A and 2B, we used IWR-1 (10 µM) and Dasatinib (50 µM) at lower concentrations for milder inhibition.

### Brightfield imaging

Embryos were embedded in 1% low-melting-point agarose to optimize positioning, covered in E3 medium supplemented with either DMSO or the different pharmacological treatments. Embryos remained embedded while time-lapse images were acquired every hour. Apparent defects in tail elongation were a physical consequence of remaining in the mounting medium while imaging. Number of somites were quantified to verify synchronous developmental staging between Control (DMSO) and treated embryos. Images were acquired using a Nikon SMZ18 stereomicroscope.

### Primary zebrafish blastula cell culture

Embryos were collected and incubated at 28°C until reaching high stage. Embryos were dechorionated and placed on agarose-coated plates containing Ringer’s medium (NaCl 5 M, KCl 1 M, HEPES 1 M, CaCl₂ 1 M). To isolate blastula cells, the yolk cell of each embryo was mechanically disrupted and removed by manual microdissection. Chemical deyolking compromises cell quality, and optimal cell viability needs to be preserved for culture. The isolated blastoderms were then collected in sterile PBS 1X, filtered through a 40 µm filter (Pluriselect) and gently dissociated into a single-cell suspension.

48-well plate were pre-treated with Matrigel to promote cell adhesion (1% v/v in PBS 1X for standard conditions or 20% v/v exclusively for experiments shown in Figure 2H). The coating was incubated for 1h at RT and then removed prior to cell seeding. Dissociated cells were collected via centrifugation at 180 g for 3 min, after which the supernatant was discarded. The pellet was resuspended in 1mL of primary cell culture medium composed by L-15 medium supplemented with 50mM HEPES pH 7.4, 5% KnockOut Serum Replacement (KSR) and 0.5% Penicillin/Streptomycin (P/S). Cells were quantified using a Neubauer hemocytometer, and approximately 30,000 cells per well were seeded. Corresponding pharmacological treatments were added to the media prior to cell seeding: Chiron (10 µM), IWR-1 (25 µM), Dasatinib (75 µM) and Rockout (50/100 µM). Plates were then transferred for live monitoring to a Zeiss Celldiscoverer 7 automated imaging system equipped with a controlled chamber to maintain the temperature at 28.5°C and 5% CO2. All cell culture procedures and tissue dissociations were performed under sterile conditions inside a class II biological safety cabinet.

### Whole-mount embryo immunostaining and phalloidin staining

Embryos were collected at 22-23hpf, when the optic cups of WT (or DMSO-treated) control embryos were folded for at least 2h. Embryos were dechorionated with pronase and fixed in 4% FA for 24h. anti-Caspase3 and anti-phospho-Histone3 immunostainings were done as in (Sousa-Ortega *et al*, 2023). Briefly, embryos were washed in PBS1x 0.5%Tx/Tw at RT for 2h, treated with cold acetone at −20°C for 20 min and incubated in freshly prepared blocking solution (10% NGS in PBS1x 0.5%Tx/Tw). Primary antibodies anti-active caspase-3 antibody (BD Biosciences, 559565) and anti-phospho-Histone H3 (Ser10) antibody (Millipore 06-570) were diluted 1:500 in blocking solution and embryos were incubated o/n at 4 °C. Embryos were washed for 6h in PBS1x 0.5%Tx/Tw, incubated again in freshly prepared blocking solution and finally incubated O/N at 4 °C in the dark with the Alexa Fluor TM 555 Goat anti-rabbit antibody. Embryos were washed in PBS1x 0.5%Tx/Tw and counterstained with DAPI (Merck) O/N at 4 °C with DAPI (Sigma) diluted 1:1000 in PBS1x 0.5%Tx/Tw.

For phalloidin staining, fixed embryos at the same stage were incubated in PBS1X 0.1% Tw with 1% v/v DMSO, phalloidin 488 (Merck) diluted at 1:200 and DAPI (Merck) diluted 1:1000.

### Whole-mount in situ hybridization

cDNA from zebrafish at stage 24hpf was used to amplify fragments of the coding sequence of tfec and wnt2 genes. The used primer were:

tfec Forward primer, 5’ TATAAAGACCGGACGGGGACAAC 3’
tfec Reverse primer, 5’ TCCAGCTACGAATCCAGGAGCTG 3’
wnt2 Forward primer, 5’ ATGAACTTTTTGCCAAATGGAA 3’
wnt2 Reverse primer, 5’ ACACCTGCAAAACCCAGTCCTGA 3’

PCR products were clones into pSCA Strataclone plasmids (Agilent). Probes were synthesized using digoxigenin-11-UTP nucleotides (ThermoFisher) and the T3 or the T7 polymerase (ThermoFisher)) depending on the insert orientation. Probes generated were used in a final concentration of a 3 ng/μl. Embryos were fixed in 4% FA at stage 23-24hpf, dehydrated in methanol, and stored at −20 °C. ISH and FISH was performed following a previous protocol in (Sousa-Ortega *et al*, 2023).

### Confocal imaging and processing

For imaging, fixed or live embryos were embedded in 1% low-melting-point agarose and mounted in FluoroDish 35mm plates. Confocal laser scanning microscopy was performed using a Zeiss LSM880 microscope. Images were processed using ImageJ2 version 2.14.0/1.54f. For in vivo imaging, live embryos were mounted at images were acquired every 8-12 min. For cell roundness quantification, we selected RPE cells from the same central region of the domain avoiding the RIM regions. The membrane of the cells was determined using the “Freehand selection” tool and geometrical properties of the cell were obtained using the “Measure” tool. For quantification of proliferative cells, cells were selected and quantified using the “Multi-point” and “Measure” tools.

### Laser Ablation and Spinning Disk Confocal Microscopy

Embryos were mounted in 35 mm glass-bottom Fluorodish dishes, oriented face-down so that the developing eyes were directly adjacent to the coverslip. After allowing the agarose to polymerize for 2 minutes, the dishes were flooded with E3 medium containing the corresponding drug. Image acquisition and laser ablation were performed using a Roper Scientific-Photometrics Spinning Disk confocal microscope. Embryos were initially located using a 10x objective, and a 40x water-immersion objective was utilized for high-resolution imaging and targeted ablation. GFP and RFP fluorescence were excited using 491 nm and 561 nm lasers, respectively.

Ablation was targeted to the RPE using a 355 nm pulsed UV photoablation laser. The ablation target was defined as a horizontal line with a thickness of 1 pixel (corresponding to 0.33 µm) drawn either perpendicular to the actomyosin cable or along the RPE cell border. Ablation was executed using 5 laser pulses at 80% laser power, with digital gain and exposure settings maintained at approximately 100 (adjusted slightly depending on embryo depth and fluorophore brightness). Time-lapse imaging was performed at a single focal plane. Images were captured continuously for 6 seconds prior to ablation and 150 seconds post-ablation, with an actual acquisition frame rate of 1 frame per 0.729 seconds.

### Recoil Velocity Quantification

Quantitative analysis of the initial recoil velocity was performed using the PyJAMAS software suite (Fernandez-Gonzalez *et al*, 2022). The analysis parameters were calibrated to the specific settings of the Roper Scientific-Photometrics microscope as follows: temporal resolution (t_res) = 0.729 s, ablation time = 0.729 s, spatial resolution (xy_res) = 0.33 µm, and the index of time zero = 13.

### CRISPR-KO mutant generation

For sgRNAs design, CRISPRScan software was used (Moreno-Mateos *et al*, 2015). The 4 sgRNAs with the higher score flanking a region of about 200bp were selected. To later evaluate sgRNA efficiency, the flanking region was subsequently amplified by PCR.

Embryo microinjections were performed at one-cell stage using a M400 Narishige microinjector with the injection volume calibrated using a calibration device. The final concentration of Cas9 mRNA was 150pg, and 4uM of each sgRNA mRNA. At 24h post-injection, genomic DNA was extracted from 10 embryos, and the PCR product of the flanking region was sequenced to verify the targeted deletion. We confirmed a 16bp deletion in the second exon that introduced a premature stop codon. To establish the stable CRISPR/KO line, F0 founders were outcrossed with WT fish, and the resulting F1 offspring were sequenced to identify carriers of the deletion. Heterozygous mutants were identified by fin-clipping and genotyping of the F1 generation. We isolated homozygous mutants from the subsequent F2 generation to maintain a stable homozygous line.

### RNAseq

#### RNA extraction, library preparation and sequencing

Embryo heads from the respective treatments were dissected and transferred to a 1.5mL RNAse-free Eppendorfs and pooled in groups of 10 heads/treatment/replicate. Samples were immediately snap-frozen in liquid nitrogen.

RNA was extracted with the RNeasy Mini Kit from QIAGEN (Cat no. 74104) following the manufacturer instructions. Extracted RNA eluted in 15uL Nuclease-free H2O was sent for library preparation and sequencing to Novogene. Poly(A) capture and reverse transcription were performed to generate cDNA libraries, followed by rigorous quality control assessments. Sequencing was carried out on an Illumina NovaSeq X platform using a 150 bp paired-end strategy, yielding approximately 12 Gb of raw data per sample.

#### Bioinformatic analysis

Reads were pre-processed trimming the low quality (lower than 20) bases at the beginning and end of the sequence. Those bp with an average quality lower than 20 were filtered out (-r --cut_right_window_size 3 --cut_right_mean_quality 20 -M 30 -W 1 -q 30 -e 30 -l 20 -w 20). Processed reads were then mapped against the last version of the zebra fish genome (danRer11) using Kallisto v0.50.1. The subsequent analysis was performed using Rstudio v.4.4.0. Genes with less than 10 reads on average were discarded (rowSums(counts(ddsTxi) >= 10)). Differential gene expression analysis was carried out using the R package DESeq2 (padj <0.05; logFC = +/−1.5; https://www.r-project.org/). GO Terms were analyzed using clusterProfiler v.4.12.6. Heatmaps were plotted with pheatmap v.1.0.13. For PCAs, volcanos, heatmaps and box plots, R package ggplot2 v.4.0.2. was used.

#### Data availability

RNAseq datasets are available in the Annotare repository from EMBL-EBI in the following accession: https://urldefense.com/v3/__http://helpdesk.ebi.ac.uk/Ticket/Display.html?id=876010__;!!D9dNQwwGXtA!RXD39PUsBt1qz8wxav45zxSKhythpiQQgEwn3A7Bf45hHxY00jSLhR6lMkszO7Vw8iG1xb0I92ra9eQ$

## Supplementary Figures

**Supplementary Figure 1.**
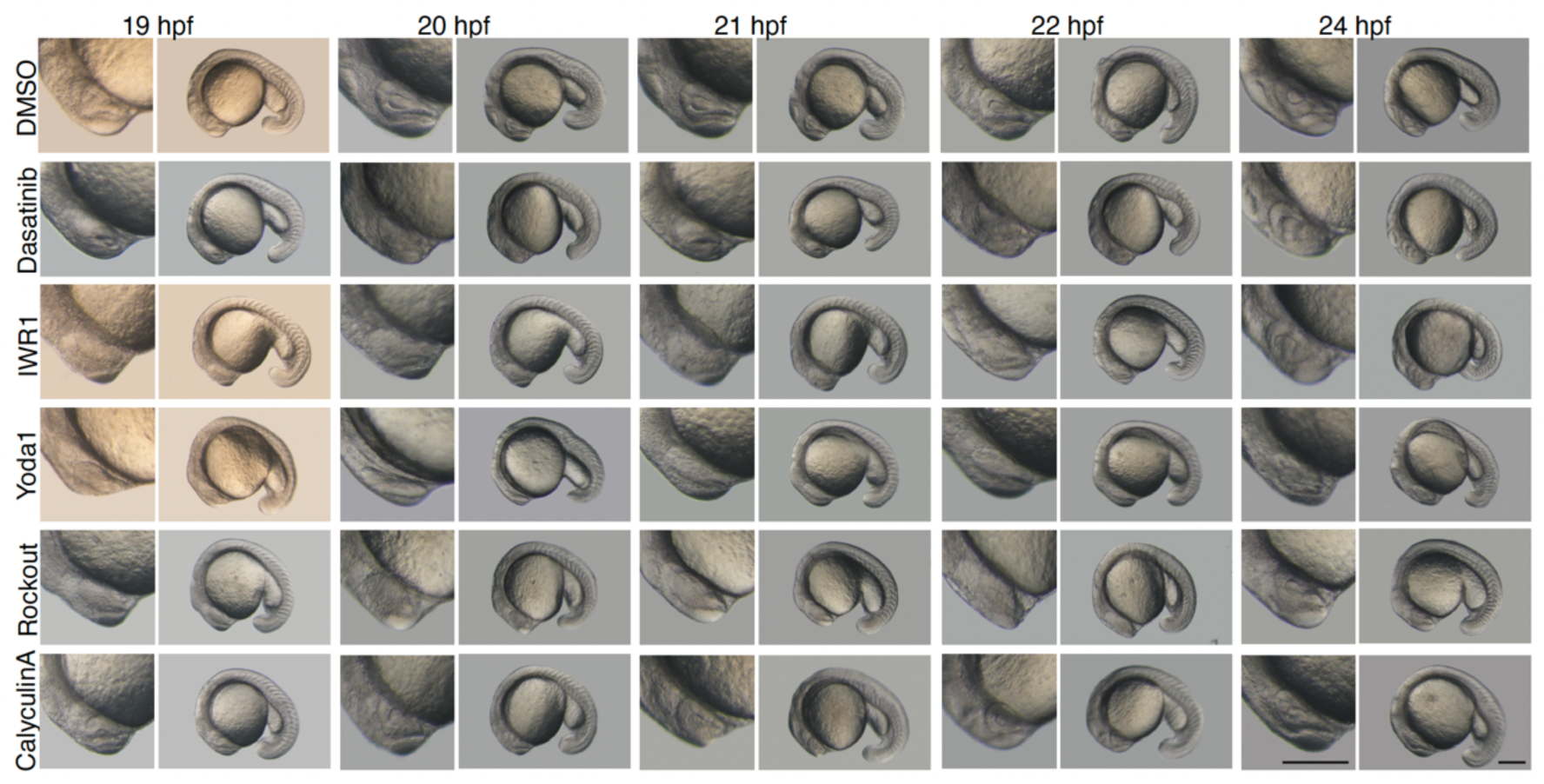
Kinetics delay of optic cup folding after signaling and cellular tension perturbations. Brightfield time-lapse images showing the global embryo development (right columns) and magnification of the corresponding optic cups (left columns) of zebrafish embryos from 19 to 24hpf. Embryos were embedded in 1% low-melting-point agarose and imaged every hour. Embryos were treated with Dasatinib (YAP inhibitor), IWR1 (Wnt/ β-catenin inhibitor), Yoda1 (Piezo1 agonist), Rockout (ROCK and actomyosin inhibitor) or Calyculin A (actomyosin hyperactivator). Optic vesicles of control (DMSO) and Calyculin-treated embryos begin to fold at 19 hpf, whereas treatments with Dasatinib, IWR1, Yoda1, and Rockout result in a consistent delay of approximately 5 hours in the onset of optic cup folding. Scale bars: 200 µm.

**Supplementary Figure 2.**
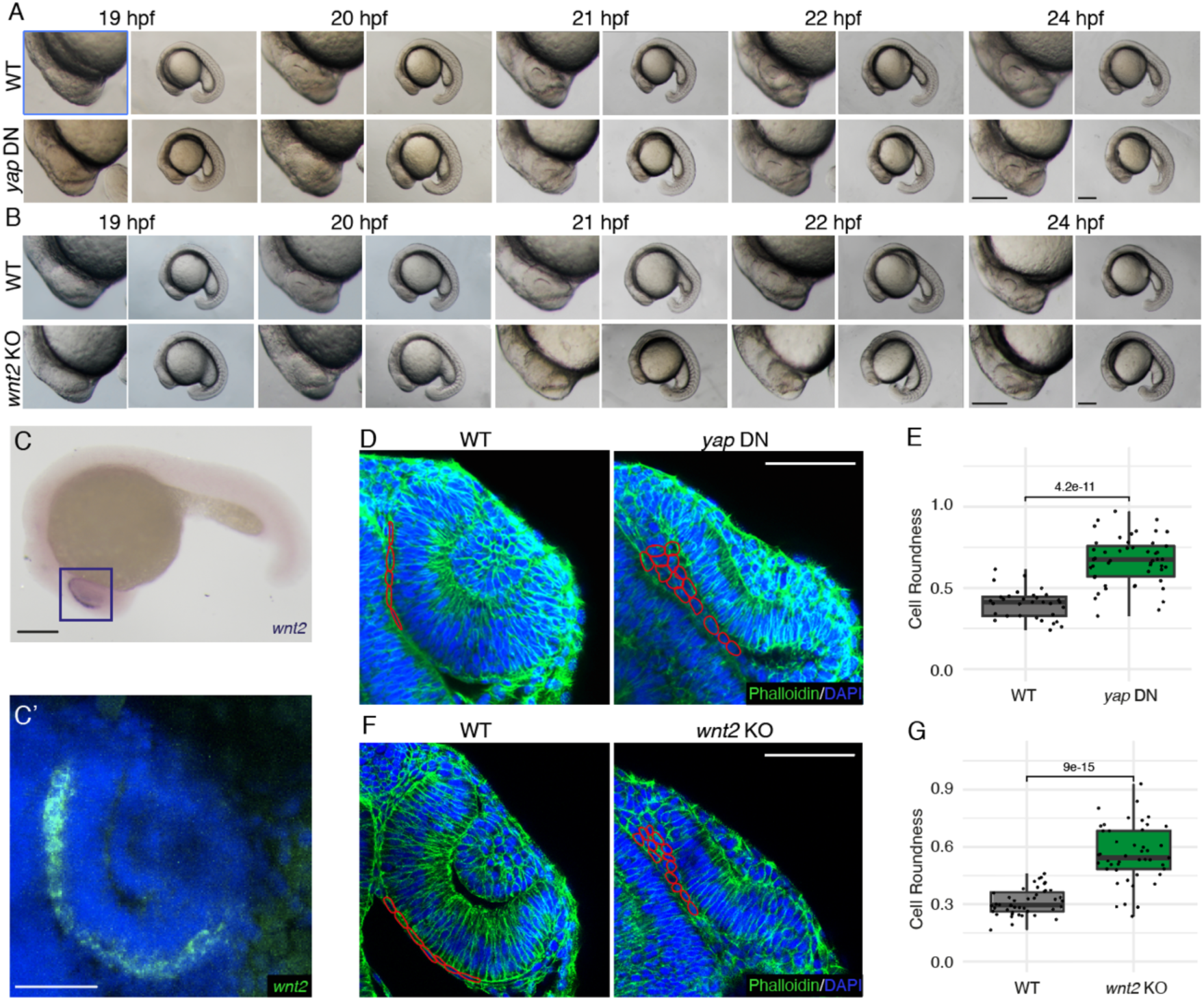
Alternative methods of inhibiting YAP and Wnt/ β-catenin corroborate their roles in regulating RPE cell geometries and ensuring a timely optic cup folding. **(A)** Brightfield time-lapse images showing the global embryo development (right columns) and magnification of the corresponding optic cups (left columns) of zebrafish embryos from 19 to 24hpf. Embryos were embedded in 1% low-melting-point agarose and imaged every hour. Embryos injected with dominant-negative forms of YAP (*yap* DN) show a consistent delay in optic cup folding of approximately 3h compared to WT siblings. Scale bars: 150 µm. **(B)** Brightfield time-lapse images showing the global embryo development (right columns) and magnification of the corresponding optic cups (left columns) of zebrafish embryos from 19 to 24hpf. Embryos were embedded in 1% low-melting-point agarose and imaged every hour. *wnt2* knockout (*wnt2* KO) embryos display a consistent delay in optic cup folding of approximately 5h. Scale bars: 150 µm. **(C)** Whole-mount in situ hybridization shows the restricted expression of *wnt2* at the RPE. Scale bar: 150 µm. **(C’)** Confocal image of the region of the eye displaying wnt2 fluorescent *in* situ hybridization (green) counterstained with DAPI (blue) confirming the restricted expression of *wnt2* in the RPE domain. Scale bar: 50 µm. **(D)** Confocal images of WNT and *yap* DN optic cups at 24hpf stage stained with Phalloidin (green) and DAPI (blue). In red, representative RPE geometries are highlighted. Scale bars: 50 µm. **(E)** Quantification of RPE cell roundness in WT and *yap* DN embryos. Box plots represent the distribution of cell roundness values; individual data points represent single cells. Data normality was assessed via the Shapiro-Wilk test. Because all the groups exhibited a normal distribution, statistical significance was determined using the parametric t-test. n=31 WT cells from 6 embryos; n=48 *yap* DN cells from 8 embryos. **(F)** Confocal images of WNT and *wnt2* KO optic cups at 24hpf stage stained with Phalloidin (green) and DAPI (blue). In red, representative RPE geometries are highlighted. Scale bars: 50 µm. **(G)** Quantification of RPE cell roundness in WT and *wnt2* KO embryos. Box plots represent the distribution of cell roundness values; individual data points represent single cells. Data normality was assessed via the Shapiro-Wilk test. Because all the groups exhibited a normal distribution, statistical significance was determined using the parametric t-test. n=44 WT cells from 6 embryos; n=45 *yap* DN cells from 6 embryos.

**Supplementary Figure 3.**
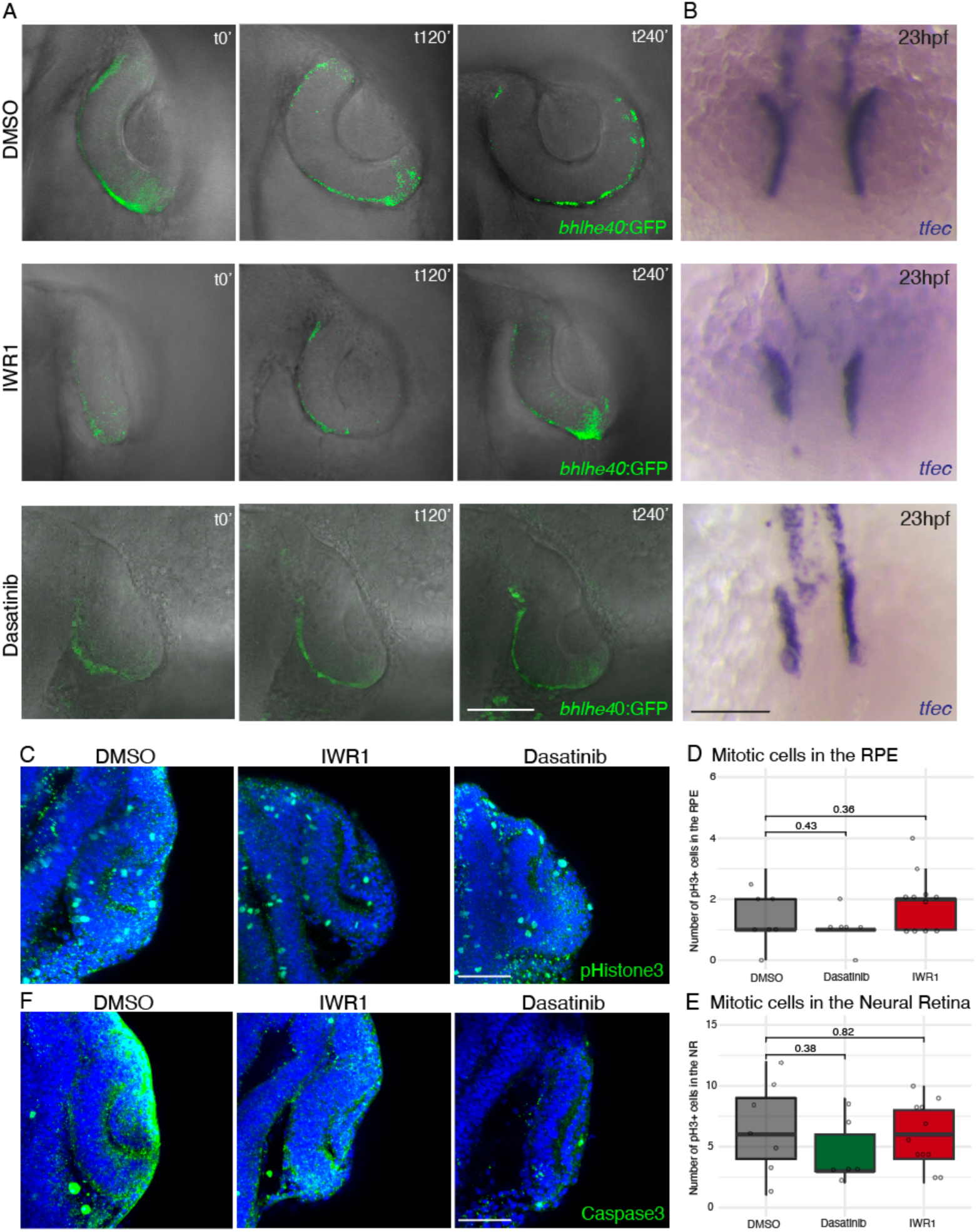
Inhibition of YAP and Wnt/ β-catenin does not affect early RPE identity specification, proliferation, and cell survival in the optic cup. **(A)** Live time-lapse imaging snapshots showing the activation of Bhlhe40, an early RPE specification marker via the reporter line *bhlhe40:*GFP (Moreno-Mármol *et al*, 2021) (green) in DMSO- (Control), and IWR1-, and Dasatinib (YAP inhibitor)-treated (β-catenin inhibitor) embryos. Time points (t) are indicated in minutes. Note that t0 correspond to the initiation of time-lapse imaging for each respective movie, which was stage 20hpf for the three different treatments. **(B)** Whole-mount *in situ* hybridization showing the expression of another early RPE specification marker *tfec* (blue) at 23 hpf in DMSO- (Control), IWR1-, and Dasatinib-treated embryos. RPE specification markers are active even though the morphogenetic folding of the optic cup is altered following YAP and Wnt/β-catenin inhibition. **(C-E)** Analysis of mitotic cells in the optic cup following IWR1 and Dasatinib treatment. **(C)** Confocal sections of representative optic cups at 24hpf stage stained for the mitotic marker phospho-Histone H3 (pHistone3, green) and counterstained with DAPI (blue) in the different treatments. **(D)** Quantification of the number of pH3+ cells in the RPE domain. No statistically significant differences were observed between control and treated groups. Box plots represent the distribution of number of pH3+ cells/RPE/embryo; individual data points represent single embryos. Data normality was assessed via the Shapiro-Wilk test. Because at least one group exhibited a non-normal distribution, statistical significance was determined using the non-parametric Wilcoxon test. n=7 WT embryos; n=6 Dasatinib-treated embryos; n=11 IWR1-treated embryos. **(E)** Quantification of the number of pH3+ cells in the NR domain. No statistically significant differences were observed between control and treated groups. Box plots represent the distribution of number of pH3+ cells/NR/embryo; individual data points represent single embryos. Data normality was assessed via the Shapiro-Wilk test. Because at least one group exhibited a non-normal distribution, statistical significance was determined using the non-parametric Wilcoxon test. n=7 WT embryos; n=6 Dasatinib-treated embryos; n=11 IWR1-treated embryos. **(F)** Confocal images of representative optic cups at 24hpf stage stained for the apoptotic marker activated Caspase-3 (Caspase3, green) counterstained with DAPI (blue) across the different treatments. Apart from nonspecific background signal, no significant cell death was detected in the RPE or NR of either control and treated embryos. Scale bars: 50 µm.

**Supplementary Figure 4.**
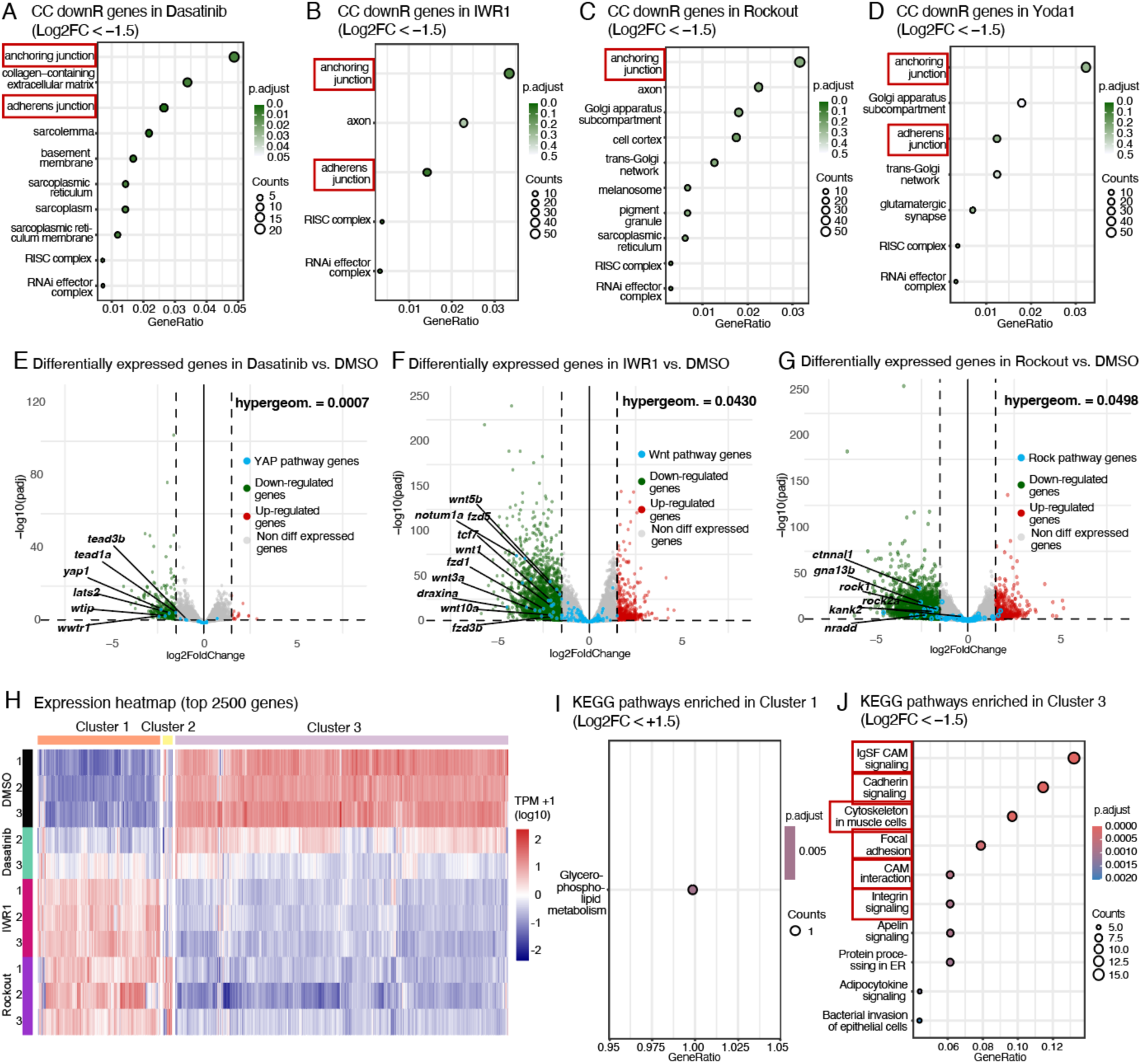
Cellular component enrichment, targeted pathway profiling and cluster analysis of common up-regulated and down-regulated genes following reduction of intracellular tension and YAP/β-catenin signaling. **(A-D)** Cellular Component (CC) GO term enrichment dot plots of down-regulated genes (with Log_2_ FoldChange ≤ −1.5) in embryos treated with Dasatinib, IWR1, Rockout, or Yoda compared to DMSO controls. The x-axis indicates gene ratio, dot size represents the gene count within each pathway, and dot color indicates the adjusted p-value. Structural terms like “anchoring junction” and “adherens junction” are strongly enriched across treatments and are highlighted by colored boxes. **(E-G)** Volcano plots showing differentially expressed genes (DEGs) in Dasatinib vs. DMSO **(E)**, IWR1 vs. DMSO **(F)**, and Rockout vs. DMSO **(G)**. y-axis represents the negative log_10_ of the adjusted p-values and x-axis represents log_2_ FoldChange. Down-regulated genes (green), up-regulated genes (red), non-differentially expressed genes (grey), and specific pathway-related genes (blue) are shown. Statistical significance for the enrichment of pathway-specific genes is indicated by the hypergeometric test (top right of each plot). Key pathway-specific down-regulated genes are labeled. **(H)** Expression heatmap of the top 2500 DEGs across DMSO, Dasatinib, IWR1, and Rockout replicates (TPM+1 log_10_ scale). Hierarchical clustering separates genes into distinct profiles, predominantly Cluster 1 (up-regulated genes in Dasatinib, IWR1, and Rockout treatments compared to DMSO), and Cluster 3 (down-regulated genes in Dasatinib, IWR1, and Rockout treatments compared to DMSO). **(I, J)** KEGG pathway enrichment plots of genes grouped in Cluster 1 (up-regulated DEGs) (with Log_2_ FoldChange ≤ +1.5) and cluster 3 (down-regulated DEGs) (with Log_2_ FoldChange ≤ −1.5). Exclusively among Cluster3 genes, we can find strong enrichment of mechanosensitive KEGG pathways related to adhesion, cytoskeleton, and contractility pathways, which are highlighted by colored boxes (CAM=Cell Adhesion Molecules).

**Supplementary Figure 5.**
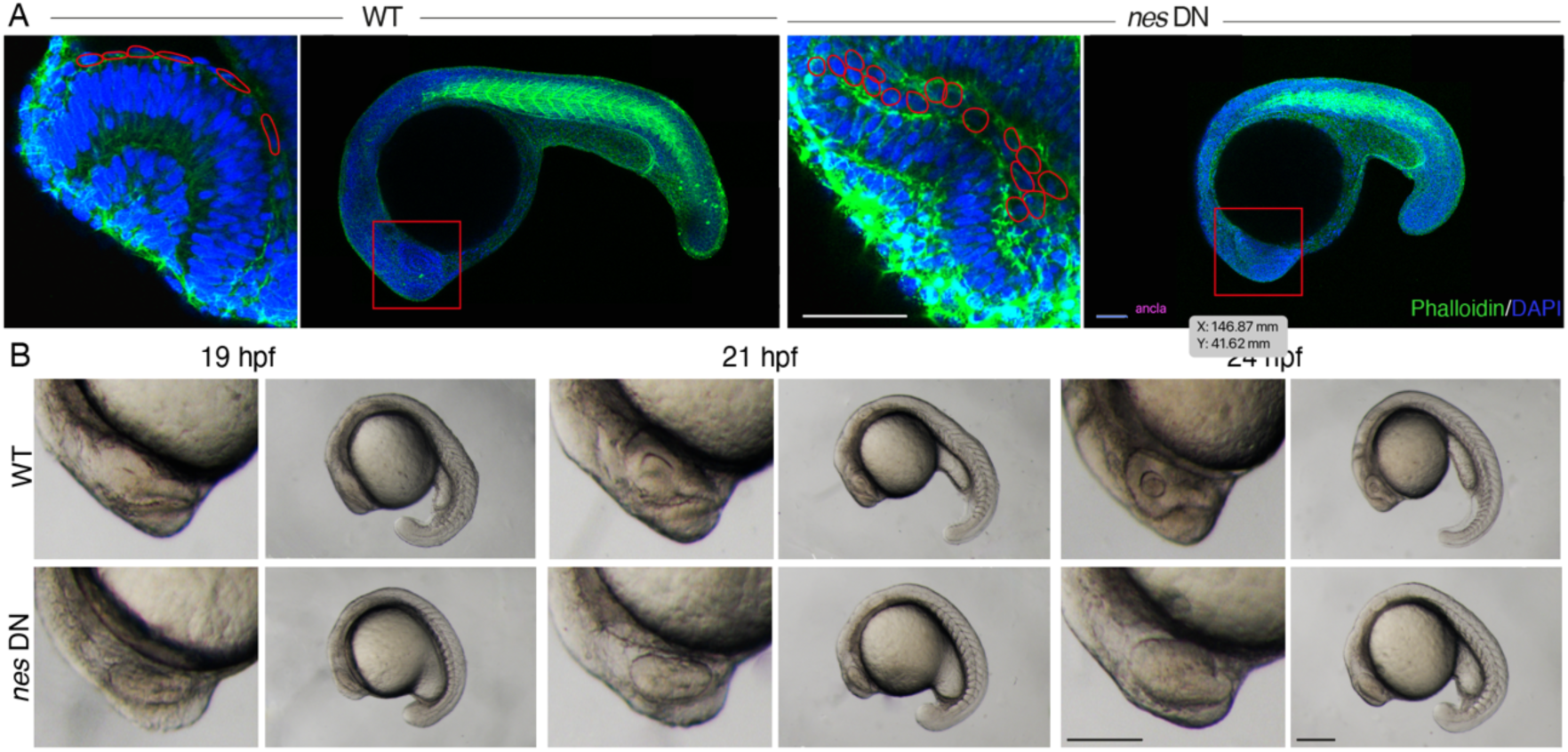
Ubiqutous inhibition of the LINC domain arrests global embryonic development rather than specifically delaying optic cup folding. **(A)** Confocal images of representative WT and *nes* DN optic cups at 24hpf stage stained with Phalloidin (green) and DAPI (blue). Whole-embryo views (right panels) and high-magnification of the eye region (left panels; indicated by red boxes). In red, representative RPE geometries are outlined. Scale bars: 50 µm. **(B)** Brightfield time-lapse images showing the global embryo development (right columns) and magnification of the corresponding optic cups (left columns) of zebrafish embryos from 19 to 24hpf. Embryos were embedded in 1% low-melting-point agarose and imaged every 2-3 hours. The delay in the onset of optic cup folding in *nes* DN embryos is accompanied by overall embryonic developmental arrest. Scale bars: 150 µm.

## Supplementary movies

**<u>Supplementary movie 1.</u>** Live time-lapse imaging of the Wnt/β-catenin activation in the RPE cells via the β-catenin reporter 7x*TCFsiam:*mCherry (*TCF:*mCherry) (red). This movie corresponds to Figure 1A. Time interval is 10 minutes per frame.

**<u>Supplementary movie 2.</u>** Live time-lapse imaging of the Wnt/β-catenin activation via the β-catenin reporter 7x*TCFsiam:*mCherry (*TCF:*mCherry) (red) and the activation of Bhlhe40, an early RPE specification marker via the reporter line *bhlhe40:*GFP (green). β-catenin activity and Bhlhe40 expression specifically co-localizes in the flattening RPE cells. Time interval is 10 minutes per frame.

**<u>Supplementary movie 3.</u>** Live time-lapse imaging of YAP and β-catenin activation in the RPE cells via the double reporter line for YAP 4X*GTIIC:*Kaede (*TEAD:*Kaede) (green) and β-catenin 7x*TCFsiam:*mCherry (*TCF:*mCherry) (red). YAP and β-catenin activity are co-localized and restricted to the flattening RPE domain. This movie corresponds to Figure 1B. Time interval is 10 minutes per frame.

**<u>Supplementary movie 4.</u>** Live time-lapse imaging of the optic cup from a WT embryo injected with *Utrophin*:GFP (actin filament contractility marker; green) and *Lyn-*dTomato (cellular membrane marker; red) mRNA. This WT embryo is the sibling control for the subsequent movie 5. Time interval is 10 minutes per frame.

**<u>Supplementary movie 5.</u>** Live time-lapse imaging of the optic cup from a *YapDN* embryo injected with *Utrophin*:GFP (actin filament contractility marker; green) and *Lyn-*dTomato (cellular membrane marker; red) mRNA. Time interval is 10 minutes per frame.

**<u>Supplementary movie 6.</u>** Live time-lapse imaging of the optic cup from a WT embryo injected with *Utrophin*:GFP (actin filament contractility marker; green) and *Lyn-*dTomato (cellular membrane marker; red) mRNA. This WT embryo is the sibling control for the subsequent movie 7. Time interval is 10 minutes per frame.

**<u>Supplementary movie 7.</u>** Live time-lapse imaging of the optic cup from a *wnt2* KO embryo injected with *Utrophin*:GFP (actin filament contractility marker; green) and *Lyn-*dTomato (cellular membrane marker; red) mRNA. Time interval is 10 minutes per frame.

**<u>Supplementary movie 8.</u>** Live time-lapse imaging of the YAP activation via the YAP reporter 4X*GTIIC:*Kaede (*TEAD:*Kaede) (green) in a DMSO-treated (Control) embryo. This movie corresponds to Figure 2A. Time interval is 10 minutes per frame.

**<u>Supplementary movie 9.</u>** Live time-lapse imaging of the YAP activation via the YAP reporter 4X*GTIIC:*Kaede (*TEAD:*Kaede) (green) in a Dasatinib-treated (YAP inhibitor) embryo. Even when YAP signaling is just mildly inhibited using low dosis of Dasatininb, the YAP reporter *TEAD*:Kaede does not show activity in the RPE, validating the pharmacological treatment. This movie corresponds to Figure 2A. Time interval is 10 minutes per frame.

**<u>Supplementary movie 10.</u>** Live time-lapse imaging of the Wnt/β-catenin activation via the β-catenin reporter 7x*TCFsiam:*mCherry (*TCF:*mCherry) (red) in a DMSO-treated (Control) embryo. This movie corresponds to Figure 2B. Time interval is 10 minutes per frame.

**<u>Supplementary movie 11.</u>** Live time-lapse imaging of the Wnt/β-catenin activation via the β-catenin reporter 7x*TCFsiam:*mCherry (*TCF:*mCherry) (red) in a IWR1-treated (β-catenin inhibitor) embryo. Even when β-catenin signaling is just mildly inhibited using low-dose of IWR1, the Wnt/β-catenin reporter *TCF:*mCherry does not show activity in the RPE, validating the pharmacological treatment. This movie corresponds to Figure 2B. Time interval is 10 minutes per frame.

**<u>Supplementary movie 12.</u>** Live time-lapse imaging of dissociated zebrafish blastula cells from reporter *TEAD:*Kaede (green) embryos and cultured in DMSO (Control). This movie corresponds to Figure 2C. Time interval is 8 minutes per frame.

**<u>Supplementary movie 13.</u>** Live time-lapse imaging of dissociated zebrafish blastula cells from reporter *TEAD:*Kaede (green) embryos injected with the focal adhesion marker *Paxillin:*mKate (red) and cultured in DMSO (Control). Yap-active cell colonies are surrounded by focal adhesions. This movie corresponds to Figure 2C. Time interval is 8 minutes per frame.

**<u>Supplementary movie 14.</u>** Live time-lapse imaging of dissociated zebrafish blastula cells from reporter *TEAD:*Kaede (green) embryos injected with the focal adhesion marker *Paxillin:*mKate (red) and cultured with Dasatinib (YAP inhibitor). Focal adhesions fail to assemble around the colonies and they do no activate YAP (*TEAD:*Kaede in green). This movie corresponds to Figure 2C. Time interval is 8 minutes per frame.

**<u>Supplementary movie 15.</u>** Live time-lapse imaging of dissociated zebrafish blastula cells from reporter *TCF:*mCherry (red) embryos cultured with DMSO (Control). This movie corresponds to Figure 2D. Time interval is 8 minutes per frame.

**<u>Supplementary movie 16.</u>** Live time-lapse imaging of dissociated zebrafish blastula cells from reporter *TCF:*mCherry (red) embryos cultured with Chiron (Wnt/β-catenin activator). Chiron boosts *TCF:*mCherry activation intensity. This movie corresponds to Figure 2D. Time interval is 8 minutes per frame.

**<u>Supplementary movie 17.</u>** Live time-lapse imaging of dissociated zebrafish blastula cells from reporter *TCF:*mCherry (red) embryos cultured with IWR1 (Wnt/β-catenin inhibitor). IWR1 suppresses *TCF*:mCherry activation. This movie corresponds to Figure 2D. Time interval is 8 minutes per frame.

**<u>Supplementary movie 18.</u>** Live time-lapse imaging of dissociated zebrafish blastula cells from reporter *TEAD:*Kaede (green) embryos cultured with DMSO (Control). This movie corresponds to Figure 2E. Time interval is 8 minutes per frame.

**<u>Supplementary movie 19.</u>** Live time-lapse imaging of dissociated zebrafish blastula cells from reporter *TEAD:*Kaede (green) embryos cultured with IWR1 (Wnt/β-catenin inhibitor). IWR1 also suppresses *TEAD:*Kaede activation. This movie corresponds to Figure 2E. Time interval is 8 minutes per frame.

**<u>Supplementary movie 20.</u>** Live time-lapse imaging of dissociated zebrafish blastula cells from reporter *TCF*:mCherry (red) embryos cultured with Chiron (Wnt/β-catenin activator). This movie corresponds to Figure 2E. Time interval is 8 minutes per frame.

**<u>Supplementary movie 21.</u>** Live time-lapse imaging of dissociated zebrafish blastula cells from reporter *TCF:*mCherry (red) embryos cultured with Chiron (Wnt/β-catenin activator) and Dasatinib (YAP inhibitor). Dasatinib suppresses *TEAD:*Kaede activation also in combination with Chiron. This movie corresponds to Figure 2E. Time interval is 8 minutes per frame.

**<u>Supplementary movie 22.</u>** Live time-lapse imaging of dissociated zebrafish blastula cells from reporter *TEAD:*Kaede (green) embryos cultured with DMSO (Control). This movie corresponds to Figure 2G. Time interval is 8 minutes per frame.

**<u>Supplementary movie 23.</u>** Live time-lapse imaging of dissociated zebrafish blastula cells from reporter *TEAD:*Kaede (green) embryos cultured with Rockout (intracellular tension inhibitor). Rockout suppresses *TEAD:*Kaede activation. This movie corresponds to Figure 2G. Time interval is 8 minutes per frame.

**<u>Supplementary movie 24.</u>** Live time-lapse imaging of dissociated zebrafish blastula cells from reporter *TCF:*mCherry (red) embryos cultured with Chiron (Wnt/β-catenin activator). This movie corresponds to Figure 2G. Time interval is 8 minutes per frame.

**<u>Supplementary movie 25.</u>** Live time-lapse imaging of dissociated zebrafish blastula cells from reporter *TCF:*mCherry (red) embryos cultured with Chiron (Wnt/β-catenin activator) and Rockout (intracellular tension inhibitor). Rockout suppresses *TCF:*mCherry activation also in combination with Chiron. This movie corresponds to Figure 2G. Time interval is 8 minutes per frame.

**<u>Supplementary movie 26.</u>** Live time-lapse imaging of dissociated zebrafish blastula cells from reporter *TEAD:*Kaede (green) embryos cultured with DMSO (Control) in a 1% Matrigel substrate. This movie corresponds to Figure 2H. Time interval is 8 minutes per frame.

**<u>Supplementary movie 27.</u>** Live time-lapse imaging of dissociated zebrafish blastula cells from reporter *TCF:*mCherry (red) embryos cultured with DMSO (Control) in a 1% Matrigel substrate. This movie corresponds to Figure 2H. Time interval is 8 minutes per frame.

**<u>Supplementary movie 28.</u>** Live time-lapse imaging of dissociated zebrafish blastula cells from reporter *TCF:*mCherry (red) embryos cultured with Chiron (Wnt/β-catenin activator) in a 1% Matrigel substrate. This movie corresponds to Figure 2H. Time interval is 8 minutes per frame.

**<u>Supplementary movie 29.</u>** Live time-lapse imaging of dissociated zebrafish blastula cells from reporter *TEAD:*Kaede (green) embryos cultured with DMSO (Control) in a 20% Matrigel substrate. Higher concentration of Matrigel in the substrate (stiffer ECM and higher ligand concentration) results in enhanced activation of *TEAD:*Kaede reporter. This movie corresponds to Figure 2H. Time interval is 8 minutes per frame.

**<u>Supplementary movie 30.</u>** Live time-lapse imaging of dissociated zebrafish blastula cells from reporter *TCF:*mCherry (red) embryos cultured with DMSO (Control) in a 20% Matrigel substrate. This movie corresponds to Figure 2H. Time interval is 8 minutes per frame.

**<u>Supplementary movie 31.</u>** Live time-lapse imaging of dissociated zebrafish blastula cells from reporter *TCF:*mCherry (red) embryos cultured with Chiron (Wnt/β-catenin activator) in a 20% Matrigel substrate. Higher concentration of Matrigel in the substrate (stiffer ECM and higher ligand concentration) results in enhanced activation of *TCF:*mCherry reporter. This movie corresponds to Figure 2H. Time interval is 8 minutes per frame.

**<u>Supplementary movie 32.</u>** Live time-lapse imaging of laser ablation of the actomyosin cable at the RPE-NR interface of a *myosin2:*GFP (green) reporter embryo treated with DMSO (Control). This movie corresponds to Figure 3C. Time interval is 30 seconds per frame.

**<u>Supplementary movie 33.</u>** Live time-lapse imaging of laser ablation of the actomyosin cable at the RPE-NR interface of a *myosin2:*GFP (green) reporter embryo treated with Dasatinib (YAP inhibitor). This movie corresponds to Figure 3C. Time interval is 30 seconds per frame.

**<u>Supplementary movie 34.</u>** Live time-lapse imaging of laser ablation of the actomyosin cable at the RPE-NR interface of a *myosin2:*GFP (green) reporter embryo treated with IWR1 (Wnt/β-catenin inhibitor). This movie corresponds to Figure 3C. Time interval is 30 seconds per frame.

**<u>Supplementary movie 35.</u>** Live time-lapse imaging of a single RPE cell laser ablation of a *myosin2:*GFP (green) reporter embryo injected with *Lyn-*dTomato (cellular membrane marker; red) and treated with DMSO (Control). This movie corresponds to Figure 3E. Time interval is 30 seconds per frame.

**<u>Supplementary movie 36.</u>** Live time-lapse imaging of a single RPE cell laser ablation of a *myosin2:*GFP (green) reporter embryo injected with *Lyn-*dTomato (cellular membrane marker; red) and treated with Dasatinib (YAP inhibitor). This movie corresponds to Figure 3E. Time interval is 30 seconds per frame.

**<u>Supplementary movie 37.</u>** Live time-lapse imaging of a single RPE cell laser ablation of a *myosin2:*GFP (green) reporter embryo injected with *Lyn-*dTomato (cellular membrane marker; red) and treated with IWR1 (Wnt/β-catenin inhibitor). This movie corresponds to Figure 3E. Time interval is 30 seconds per frame.

